# The cellular Notch1 Protein Promotes KSHV reactivation in an Rta-dependent manner

**DOI:** 10.1101/2022.10.21.513206

**Authors:** Jennifer DeCotiis-Mauro, Sun M. Han, Helena Mello, Corey Goyeneche, Giuseppina Marchesini-Tovar, Lianhua Jin, Vivian Bellofatto, David M. Lukac

**Affiliations:** Dept of Microbiology, Biochemistry, and Molecular Genetics, Rutgers Biomedical and Health Sciences, New Jersey Medical School, Rutgers University, Newark, NJ; School of Graduate Studies, Rutgers Biomedical and Health Sciences, Health Science Campus at Newark, Rutgers University, Newark, NJ

**Author notes:** authors contributed equally to this work (Jennifer DeCotiis-Mauro, Sun M. Han).

## Abstract

The cellular Notch signal transduction pathway is intimately associated with infections by Kaposi’s sarcoma-associated herpesvirus (KSHV) and other gamma-herpesviruses. RBP-Jk, the cellular DNA binding component of the canonical Notch pathway, is the key Notch downstream effector protein in virus-infected and uninfected animal cells. Reactivation of KSHV from latency requires the viral lytic switch protein, Rta, to form complexes with RBP-Jk on numerous sites within the viral DNA. Constitutive Notch activity is essential for KSHV pathophysiology in models of Kaposi’s sarcoma (KS) and Primary Effusion Lymphoma (PEL), and we demonstrate that Notch1 is also constitutively active in infected Vero cells. Although the KSHV genome contains >100 RBP-Jk DNA motifs, we show that none of the four isoforms of activated Notch can productively reactivate the virus from latency in a highly quantitative trans-complementing reporter virus system. Nevertheless, Notch contributed positively to reactivation because broad inhibition of Notch1-4 with gamma secretase inhibitor (GSI) or expression of dominant negative mastermind-like1 (dnMAML1) coactivators severely reduced production of infectious KSHV from Vero cells. Reduction of KSHV production is associated with gene specific reduction of viral transcription in both Vero and PEL cells. Specific inhibition of Notch1 by siRNA partially reduces production of infectious KSHV, and NICD1 forms promoter-specific complexes with viral DNA during reactivation. We conclude that constitutive Notch activity is required for robust production of infectious KSHV, and our results implicate activated Notch1 as a pro-viral member of a MAML1/RBP-Jk/DNA complex during viral reactivation.

**Importance:** Kaposi’s sarcoma-associated herpesvirus (KSHV) manipulates the host cell oncogenic Notch signaling pathway for viral reactivation from latency and cell pathogenesis. KSHV reactivation requires that the viral protein Rta functionally interacts with RBP-Jk, the DNA binding component of the Notch pathway, and with promoter DNA to drive transcription of productive cycle genes. We show that the Notch pathway is constitutively active during KSHV reactivation and is essential for robust production of infectious virus progeny. Inhibiting Notch during reactivation reduces expression of specific viral genes yet does not affect growth of the host cells. Although Notch cannot reactivate KSHV alone, the requisite expression of Rta reveals a previously unappreciated role for Notch in reactivation. We propose that activated Notch cooperates with Rta in a promoter-specific manner that is partially programmed by Rta’s ability to redistribute RBP-Jk DNA binding to the virus during reactivation.

## Introduction

Human gammaherpesviral infections and pathogenesis are dependent on interactions of the virus with the cellular Notch signal transduction pathway (1). In uninfected cells, the canonical model of Notch signaling involves activation of the Notch protein by proteolytic release of the Notch Intracellular Domain (NICD)1 from the membrane-associated Notch1 receptor (2, 3). NICD1 then translocates to the cell nucleus, binds to DNA-bound Recombination signal binding protein (RBP)-Jk and activates transcription by displacing a transcriptional co-repressor complex from RBP-Jk and assembling a transcriptional co-activator complex.

Reactivation of the gammaherpesvirus Kaposi’s sarcoma-associated herpesvirus (KSHV) requires the KSHV Replication and transcriptional activator (Rta) to assemble a complex with RBP-Jk and DNA to activate transcription of essential viral genes (4–27). Reflecting the central roles of Rta and RBP-Jk in reactivation, the KSHV genome contains greater than 100 consensus RBP-Jk DNA motifs, and Rta and RBP-Jk co-occupy many viral promoters during reactivation in B cells (28). However, although both Rta and NICD1 target RBP-Jk, we and others have shown that only Rta is sufficient to productively reactivate KSHV in B cell models of latency (6, 29). Indeed, our recent work showed that ectopic expression of Rta in infected Vero cells induced profound production of infectious virus assayed by a robust and sensitive trans-complementing reporter virus-cell system (30). In parallel experiments, this highly sensitive assay revealed that ectopic expression of NICD1 failed to produce detectable virus (30). Thus, this highly sensitive Vero-cell reactivation system confirmed conclusions from human B cell studies and is ideal to compare transcriptional mechanisms of Rta and Notch because its robustness exceeds those of other infection models.

Despite ectopic NICD1’s insufficiency to reactivate KSHV alone, the literature suggests that Notch signaling contributes to regulation of KSHV infection. Notch can activate viral productive cycle genes without inducing complete reactivation in B and endothelial cell models of latency (29, 31). In infected primary effusion lymphoma (PEL) cells, conditionally-expressed NICD1 activated a subset of viral genes that were different than those activated by Rta (29). Nevertheless, a broader review of the literature shows that our understanding of Notch’s role in KSHV reactivation remains debatable. One study concluded that ectopic NICD1 is capable of inducing complete reactivation (32) by directly activating Rta’s promoter and a second study indicated that Notch inhibits reactivation in neighboring cells by indirectly repressing Rta expression (33). The interpretation of experiments intended to analyze the role of Notch signaling in KSHV reactivation are complicated by two observations: first, Notch signaling is constitutively-active in all KSHV-infected cells, and second, one or more of the activated Notch proteins (i.e. NICD1-4) is required for growth and survival of KSHV-infected cells (31, 34–40). Indeed, Notch isoforms 1, 2, and 4, Notch activating ligands, and numerous Notch target genes are overexpressed and active in KS lesional tissue, PEL cells, and in KSHV-infected endothelial cells.

Notch signaling is also reciprocally impacted by KSHV infection in myriad and complex ways. The expression of Notch ligands and receptors are increased by specific latent and lytic viral proteins, including Rta, in infected lymphatic endothelial cells (31, 35, 37–39). Notch signaling is reprogramed when the viral immediate early protein Rta stimulates RBP-Jk DNA binding and by proteolytic processing of Notch effector proteins (6, 9, 28, 41). Rta-dependent stimulation of RBP-Jk binding to the KSHV Mta promoter converts it to a Notch-responsive gene (6). Moreover, our ChIP/Seq studies demonstrated that RBP-Jk binding to the viral genome is dramatically redistributed during reactivation, but Rta and RBP-Jk do not co-occupy every newly identified RBP-Jk site (28). Taken together, these data suggest that Notch plays an essential and complementary role with Rta in KSHV reactivation. While it is clear that Rta is the only viral protein that is necessary and sufficient to reactivate the complete KSHV lytic cycle, a complete understanding of Rta’s reactivation function must consider the intracellular environment that is replete with potent, constitutive Notch activity.

In the present study, we use the KSHV-infected Vero cell model and multiple complementary methodologies to demonstrate that Notch signaling is key to viral reactivation. Specifically, we demonstrate that exposing infected cells to the Notch small-molecule-inhibitor DAPT, or to Notch-inhibiting dominant negative Mastermind-like (dnMAML)-1, severely reduces production of infectious KSHV, after either chemical induction of virus or forced expression of Rta. Inhibition of KSHV virion production is accompanied by reduction of selected viral transcripts and proteins. Importantly, reactivation stymied by DAPT treatment is not the result of cell growth reduction by the drug. While we show that none of the four Notch isoforms are sufficient to reactivate the virus, transfection of specific siRNAs implicates Notch 1 as one of the isoforms that contribute to Rta-induced reactivation. We observe formation of specific Notch1/DNA complexes on the viral genome during reactivation. Finally, we also observe a block in reactivation, which is measured here by a decrease in viral DNA production, by inhibiting Notch in naturally KSHV-infected PEL cells. Taken together, our study illustrates that although Rta is necessary and sufficient for KSHV reactivation, viral reactivation relies on continued endogenous Notch signaling, likely by working with Rta cooperatively during the early steps of reactivation.

## Materials and Methods

### Plasmids

All plasmids were grown in DH5α *E. coli* and purified using the cesium chloride maxi-prep method (42). All prepared plasmids were confirmed via restriction enzyme digestion and/or sequencing. pcDNA3-FLg50 expresses full length Rta protein from a genomic allele of KSHV ORF50 (43). pCAGGS:dnMAML1-GFP and pEGFP-N1-dnMAML1 express dominant negative mutants of MAML1 fused to GFP and were gifts of Dr. Signe V. Horn Pederson (44). pEGFP-N1-H2b (referred to as H2b-GFP) expresses GFP-fused histone H2b and was a gift of Dr. Geoff Wahl (Addgene plasmid #11680) (45). pcDNA3.1-HisLacZ (13) expresses β-galactosidase reporter protein under control of the HCMV enhancer. The reporter plasmids 4xCSL-luciferase (Addgene plasmid #41726) and 4xCBS-luciferase encode firefly luciferase under the control of four RBP-Jk binding sites are were gifts of Dr. Raphael Kopan and Dr. Rhett Kovall, respectively (46, 47). p3xFLAG-NICD 1-4 express Notch intracellular domains (NICD) 1-4 from the p3xFLAG-CMV-7 expression vector and were gifts of Dr. Raphael Kopan (48). pCMV-NICD1 and pCMV-NICDΔPEST express WT and PEST-domain-deleted alleles of NICD1, respectively, and were gifts of Tom Kadesch. pcDNA3.1-ORF50ΔSTADV5 expresses transcriptionally inert, dominant negative KSHV Rta (nts 1-1590) fused to the V5 epitope tag (13). pCMV-NICD1-P12 and NICD1-L1594P express NICD1 alleles containing different activating mutations found in human T cell acute lymphoblastic leukemia lymphoma cells provided by Dr. Jon C. Aster (49). pcDNA3 (Invitrogen) was used as an empty vector control (“Vec”) and DNA filler for all experiments.

### Cell Culture

293 MSR-tet OFF and Vero rKSHV.294 cells (gifts of Dr. Jeff Vieira) were maintained as previously described (30, 50). BC3 cells (a gift of Dr. Ethel Cesarman) and BCBL-1 cells were maintained in RPMI medium supplemented with 12% or 10% Fetal Bovine Serum (FBS) respectively, 1% glutamine, 50uM β-mercaptoethanol, and 1% penicillin/streptomycin.

Viral reactivation was induced chemically by treatment of cells with 1 mM Valproic acid (VPA; Sigma, P4543; prepared in dH_2_O and filtered through 0.22uM surfactant-free cellulose acetate (SFCA)). Notch activity was inhibited chemically by treatment of cells with (2*S*)-*N*-[(3,5-Difluorophenyl)acetyl]-L-alanyl-2-phenyl]glycine 1,1-dimethylethyl ester (DAPT; Selleckchem, S2215); concentrations are given in the figure legends. The solvent vehicle for DAPT was dimethyl sulfoxide (DMSO), which was used as the control, vehicle (“Vh”) treatment in all experiments.

### Transfections

Vero rKSHV.294 cells were plated at a concentration of 2×10^5^ cells per well of 6-well plates and incubated at 37°C overnight. The following day, 2.5 ug of total plasmid DNA and 5 uL of vortexed TransIT-LT1 reagent were added to 250 uL incomplete DMEM and incubated at RT for 30m. The mixture was added to the Vero rKSHV.294 cells dropwise, and cells were incubated for 48 or 72h at 37°C, for protein harvest or SeAP assay respectively. Specific plasmids used in transfections are indicated in respective figure legends.

For siRNAs, Vero rKSHV.294 cells were plated at a concentration of 2×10^5^ cells per well of six-well plates and incubated overnight at 37° C. The following day, 30 pmol of either control siRNA-A (Santa Cruz, sc-37007), Notch 1 siRNA (h) A (Santa Cruz, sc36095A), or Notch 1 siRNA (h2) (Santa Cruz, sc-44226A) were diluted in 150 uL of incomplete DMEM and mixed with 9 uL of Lipofectamine RNAiMAX reagent previously diluted in incomplete DMEM per manufacturers’ suggestions. The mixtures were incubated for 5m at RT, then added to cells.

### Secreted Alkaline Phosphatase (SeAP) Assay

Virus-containing media from Vero rKSHV.294 cells were transferred to 293 MSR-tet-OFF cells previously plated at a density of 2×10^5^ cells/well in 6-well plates 24h earlier. 293 MSR-tet-OFF cells were returned to the incubator for 72h (unless otherwise indicated) at 37°C. SeAP-containing media were harvested, and 20 or 25 uL were assayed using either the Great EscAPe SeAP kit (Takara/Clontech) or the Phospha-Light System (Applied Biosystems). Individual data points show results from duplicate, triplicate or quadruplicate technical replicates from one to ten biologic replicates in each figure.

### Protein extractions and assays

Cells were washed with 1x PBS, and lysed using RIPA buffer (150 mM NaCl, 1% NP40, 0.5% sodium deoxycholate, 0.1% SDS, 50 mM Tris pH 8.0) supplemented with protease inhibitor cocktail (Sigma P2714) and 1 mM DTT. Dishes were scraped, contents were transferred to microfuge tubes, debris was pelleted by centrifugation, and supernatants were transferred to new microfuge tubes. Total protein concentrations were determined by Protein Assay (Bradford; Bio-Rad).

For luciferase and β-galactosidase quantitation, at 48h post-transfection, cells were washed twice with 1x PBS, then lysed in 125 uL 1x reporter lysis buffer (Promega) for 20min at RT. Dishes were scraped, contents were transferred to microfuge tubes, debris was pelleted by centrifugation, and supernatants were transferred to new microfuge tubes. Luciferase activity was measured in the BD monolight 3010 luminometer (BD Biosciences) using 20 uL of each extract into which 20 uL luciferase assay substrate was injected (Promega). β-galactosidase activity was measured by absorbance at 420 nm in the Tecan Genios plate reader using 30 uL of each extract mixed with 30 uL of 2x Assay Buffer (Promega).

### SDS-PAGE and Western Blotting

Equal amounts of protein, as determined by Bradford Assays, were mixed with 6x Laemmli buffer containing freshly added β-mercaptoethanol, heated at 95°C for 5m, then loaded on gels. Proteins were transferred via electroblotting to nitrocellulose (Main Manufacturing LLC) or PVDF (GVS Filter Tech 1212783; PVDF membranes only were initially dipped in 100% methanol). Membranes were blocked with 5% milk in 1x PBS/0.1% Tween 20 (blocking buffer) for at least 30m, then washed with 1x PBS/0.1% Tween 20 and 1x PBS. Primary antibodies were diluted in blocking buffer then incubated with the blots for 4h at RT or overnight at 4°C; antibodies and dilutions were as follows: anti-Notch1 (1:200, Santa Cruz sc-32745, or bTAN20 Developmental Studies Hybridoma Bank, deposited by Dr. Spyros Artavanis-Tsakonas), anti-Rta (1:4000, D3861 (14)), anti-FLAG (1:1000, M2 Sigma Aldrich F1804, or PA1-984B Thermo scientific), anti-GAPDH (1:1000, BioLegend 919501), anti-GFP (1:200, B-2 Santa Cruz sc-9996) anti-ORF59 (1H10; (14)), anti-KbZIP (51) or anti-histone H3 (1:4000 Proteintech 17168-1-AP), and anti-Mta (52). The membranes were then washed with 1x PBS/0.1% Tween 20 and 1x PBS and incubated for 30m with secondary antibodies diluted in blocking buffer; antibodies and dilutions were as follows: Goat anti-Mouse IgG-h+1 HRP conjugated (1:5000, Bethyl Labs A90-116P), Rabbit TruBlot anti-Rabbit IgG (1:1000, Rockland 18-8816-33), Goat anti-Rabbit IgG-h+1 HRP conjugated (1:5000, Bethyl A120-101P). Membranes were partially dried and incubated for up to 2m in Pierce ECL Western Blotting Substrate (Thermo Scientific, 32016). Signals were detected by exposure to autoradiography film (Denville, E3012) for up to 24h, or digitally captured using myECL imager (Thermo Scientific) or Chemidoc (BioRad).

### Quantitative Reverse Transcription-real-time Polymerase Chain Reaction (RT-qPCR)

Total RNA was extracted from Vero rKSHV.294 cells using the RNeasy kit (Qiagen) as per manufacturer’s suggestions. DNA contamination was removed using the TURBO DNA-free kit (Invitrogen) as per manufacturer’s suggestions. Transcripts were quantitated by qRT-PCR using the iTaq Universal SYBR Green One-Step Kit (Bio-Rad) following manufacturer’s suggestions. 5 to 100 ng. of input RNA was amplified using KSHV-specific unique coding sequence primers (53) or β-actin primers (54) in a CFX96/C1000 instrument (Bio-Rad). Reaction conditions were 10 mins 50°C for RT, 1 min 95°C, then 40 cycles of 10 sec 95°C/10 sec 60°C. Amplification specificity was confirmed for all primer pairs by melt-curve analysis and parallel control reactions that lacked template RNA or reverse transcriptase.

### vDNA quantitation

BC3 cells were treated with indicated chemicals for 65h and total DNA was purified from cell lysates using the FlexiGene DNA kit (Qiagen). Intracellular viral DNA was measured by real-time quantitative (q) PCR of the viral genome; GAPDH was the internal control. qPCR reactions were performed using SYBR Green PCR Master Mix (Thermo Scientific). Input DNA (100 ng) was amplified by incubating at 50°C for 10m and 95°C for 10m, then cycled 39 times with melting at 95°C for 15 sec and annealing at 60°C for 1m. Fold DNA was calculated using the ΔCt method and all values were normalized to that of VPA + vehicle, which was set to 100%.

### Immunofluorescence

Cells were harvested in 15 mL conical tubes, washed twice with 1x Phosphate Buffered Saline (PBS), and resuspended at a density of 1.25×10^6^ cells/mL in 1x PBS. 15-20 mm circles were drawn on glass microscope slides with a Super PAP pen (ThermoFisher 008899), then coated by a 10m incubation with 4 uL poly-L-lysine. Polylysine was removed, 200 uL of cells/circle were adhered for 30m, then fixed in pre-chilled methanol/acetone (50/50 v/v) at –20°C for 5 to 10min. Slides were air dried for 10min at RT and cells were blocked with 3% BSA/1% glycine in 1x PBS. The following primary antibodies were diluted in 1:1000 in blocking buffer and incubated for 1h with the cells: anti-Rta (D3861; (14)), anti-K8.1 (Santa Cruz Bio sc-65446, 1:1000), or anti-ORF59 (1H10; (14)). Cells were washed with 1x PBS/0.4% Tween20 and 1x PBS, then the following secondary antibodies were diluted 1:500 in blocking buffer and incubated for 1h with the cells: anti-rabbit DyLight 448 (Thermo Scientific, 35552), anti-rabbit DyLight 550 (Thermo Scientific, 84541), or anti-mouse DyLight 550 (Thermo Scientific, 84540). Cells were re-washed, and coverslips were mounted to slides using Vecta-shield with DAPI (Vector Labs).

### Cell Growth Quantitation

Vero rKSHV.294 cells were plated at a density of 6.7 x 10^3^ cells/well of a 96-well dish. 24 h later, cells were treated with 1 uL vehicle alone (control) or containing DAPT (final concentration of 1 or 5 uM), in triplicate. Three mock wells were also prepared with media only for each time point. At the post-treatment times indicated, cell media was replaced with 200 uL growth assay media (180 uL DME + 20 uL Alamar Blue (BioRad BUF012A), and incubation was continued for 4 hr at 37°C. Alamar Blue fluorescence was measured by excitation at 544 nm/emission at 590 nm in a SpectraMax M2 plate reader (Molecular Devices). Mock values were averaged and subtracted from each experimental value to yield corrected values. All corrected values were then normalized to 0h values that had been set to the relative value of 1 for each treatment.

### Chromatin Immunoprecipitation (ChIP)/Polymerase Chain Reaction (PCR)

Modified from (55, 56). Vero rKSHV.294 cells were plated at (1.5×10^6 cells per 100mm plate). 24h later, cells were transfected in triplicate with 12.5 ug total of pcDNA3 or pcDNA3-FLg50 using TransIT LT1 (Mirus) per manufacturer’s protocol. 24h after transfection, chromatin was cross-linked by replacing growth medium with 8 mL of serum-free medium containing 1.42% methanol-free formaldehyde (Peirce) and incubating for 8m at room temperature. Formaldehyde was quenched with 125 mM glycine for 5m at room temperature, then washed twice with 10 mL ice cold 1x PBS. Cells were scraped into fresh 10 mL of ice cold 1x PBS, transferred to 15 mL conical tubes, and combined with an additional 2 mL of 1x PBS that was used to rinse residual cells from each plate. Cells were pelleted by centrifugation at 2000g, 5m, at 4° C. Pellets from single plates were lysed by resuspension in 1 mL of ChIP buffer (150 mM NaCl, 50 mM Tris-HCl (pH 7.5), 5 mM EDTA, 0.5% v/v NP-40, 1.0% v/v Triton X-100), and supplemented with protease inhibitor cocktail (Sigma P2714) by repeated slow-pipetting after transfer to 1.5 mL Bioruptor Micro tubes (Diagenode) on ice. Incubation on ice was continued for 10m, and nuclear pellets were recovered by centrifugation at 12,000g for 1m at 4° C. Nuclear pellets were resuspended in 300 uL sonication buffer (ChIP buffer plus 1.0% v/v Sodium dodecyl-sulfate), and chromatin sonicated in the Bioruptor Pico (Diagenode) to average size of 250 bp (typically 15 cycles of 30s on/30s off, vortexing every 5 cycles). Nuclear lysates were as cleared by centrifugation at 12,000g, 10m, 4° C, and aliquoted in 2 million cell equivalents per subsequent ChIP. Some chromatin was set aside for analysis of input quality and quantity.

ChIPs were performed by adding anti-NICD1 antibody (Cell Signaling 3608) or equal amounts of non-specific rabbit IgG (Sigma PP64) to individual chromatin aliquots: 0.55 or 1 ug of each antibody were added to 1 million or 2 million cell equivalents, respectively. ChIPs were incubated 16-24h at 4° C with nutation. Chromatin was cleared by centrifugation at 12000g for 10 mins at 4° C and mixed with magnetic Protein A/G beads (Cytiva 17152104010150) that had been prewashed with ChIP buffer and blocked with ChIP buffer/1% v/v Bovine Serum Albumin. ChIP’d chromatin was recovered by incubating with beads for 45m at 4° C with nutation. Beads were washed 6 times with 1 mL each of ChIP buffer on a magnetic tube rack (Invitrogen). After removing the final wash, DNA was recovered by resuspending the magnetic beads in 100 uL 10% w/v Chelex 100/dH_2_0 suspension (Bio-Rad) and boiling the mixture for 10m. After cooling, proteins were removed by adding 20 ug of Proteinase K, vortexing, and incubating on a thermal mixer at 55° C for 30m at 1000 rpm. Proteinase K was inactivated by boiling samples for 10m, then centrifuged for 1m at 12000g, 4° C. 80 uLs of supernatant was transferred to new tubes. Fresh 120 uLs of dH_2_O was added to the Chelex resin, vortexed for 10s, and centrifugation repeated. 120 uL of that supernatant was removed and mixed with first 80 uL aliquot.

Purified DNA was analyzed using real-time qPCR with SYBR green (Applied Biosystems 4472918); primer sequences are available upon request. Fold enrichment of NICD1 on vDNA was calculated using the ΔCt method, comparing the Ct for anti-NICD1 to IgG first for each sample, then EV to Rta-transfected samples second. Final folds were determined by scaling to input chromatin and amount of NICD1 expression in each condition. Data shown represent results of four independent transfections performed in duplicate or triplicate, and ChIP’d twice per transfection. Two ChIPs from one transfection used alternate control non-specific IgG (Cell Signaling 2729S) or anti-GFP (Cell Signaling 2956) to confirm results.

### Statistical Analyses

Outlying data points in each experiment were identified using Peirce’s criterion and eliminated from final quantitation and graphs (57).

## Results

### Notch1 is constitutively active in KSHV-infected Vero cells

To understand the complex interactions of Notch signaling with KSHV infection, we have employed an infected Vero cell model. These cells have emerged as a robust system for understanding host and viral regulation of productive KSHV reactivation. Infected Vero cells provide a number of improvements over other reactivation models, including their ease of transfection. Gantt and colleagues (50) developed a Vero cell line that produces a recombinant KSHV reporter virus upon reactivation that is quantifiable by infecting a trans-complementing 293 cell line. Using this system, we previously demonstrated that ectopic, activated Notch1 (NICD1) was insufficient to produce infectious virus, confirming work from studies of PEL cells (6, 29, 30).

Many cellular models of KSHV infection demonstrate constitutive activation of NICD1 (31, 34, 36–38) which is required to maintain cell growth and survival (36–38). Based on the dramatic association of Notch activity with KSHV infection in those cells, we sought to determine the status of Notch1 in infected Vero cells. We prepared total protein extracts from infected Vero cells. Western blotting with anti-Notch1-specific antisera demonstrated a protein band doublet that migrated with an apparent molecular weight (MW) of 100-120 kDa (Fig. 1A). The upper band matched the migration of the band observed from control, uninfected Vero cells transfected with an NICD1 expression vector (Fig. 1A). Therefore, we conclude that Notch1 is constitutively active in latent, KSHV-infected Vero cells. Interestingly, significantly-less activated Notch1 is observed in uninfected Vero cells (center lanes of Fig 1A, quantitated in Fig 1B). These data suggest that latent infection of Vero cells reduces activation of endogenous Notch1. However, we can’t exclude the possibility that the reduction of Notch1 activation trivially results from the Puromycin selection of the virus episome in rKSHV.294 cells. Others have shown that expression of one or several productive cycle KSHV proteins could induce expression of Notch pathway components in 293 cells (39).

**Figure 1.**
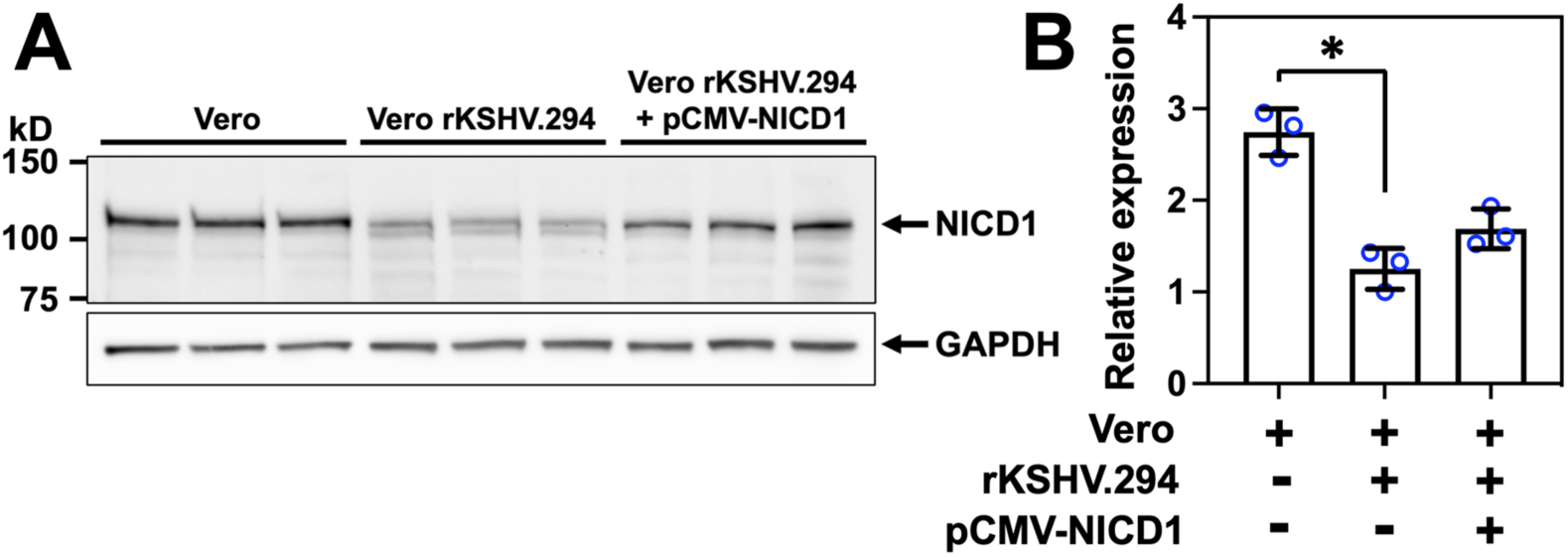
Endogenous Notch1 is constitutively active in infected Vero cells. **A**. Total cellular protein was harvested in RIPA buffer from triplicate cultures of 4 x 10^5^ cells Vero rKSHV.294 cells, uninfected Vero cells, or uninfected Vero cells transfected plasmid expressing NICD1. Equal amounts of each extract were separated on a 4-12% PAGE gradient gel electrophoresed in Tris-MOPS-SDS buffer. Proteins transferred to nitrocellulose were probed with Notch1-specific bTAN20 antibody (1:500) or anti-GAPDH antibody (1:5000), followed by HRP-conjugated anti-rat secondary antibody (1:5000). Bands were visualized by ChemiDoc (BioRad). **B. Quantitation.** NICD1 and GAPD bands were measured by ImageJ (134). NICD1 values were divided by GAPDH values and plotted. *p<0.002 by t test.

### Inhibiting Notch activation reduces KSHV reactivation

To determine whether endogenous, constitutive Notch activity contributed to KSHV reactivation, we measured the effect of DAPT treatment on mature infectious virus production from Vero rKSHV.294 cells. Cells were pre-treated with DAPT or the vehicle control (“Vh”; DMSO) for 24h, then transfected with Rta expression vector to induce viral reactivation or empty vector (Vec) control. When Notch is active, which is the case in the Vh only cultures, ectopic Rta reactivates virus to approximately 700-fold greater than Vec, as we previously published (Fig. 2A) (30). In cells where Notch is inhibited, which is the case in the DAPT-treated cells, and viral reactivation is attempted by Rta addition, reactivation is stymied. These data suggest that constitutive Notch activity is necessary for optimal viral production (Fig. 2A).

**Figure 2.**
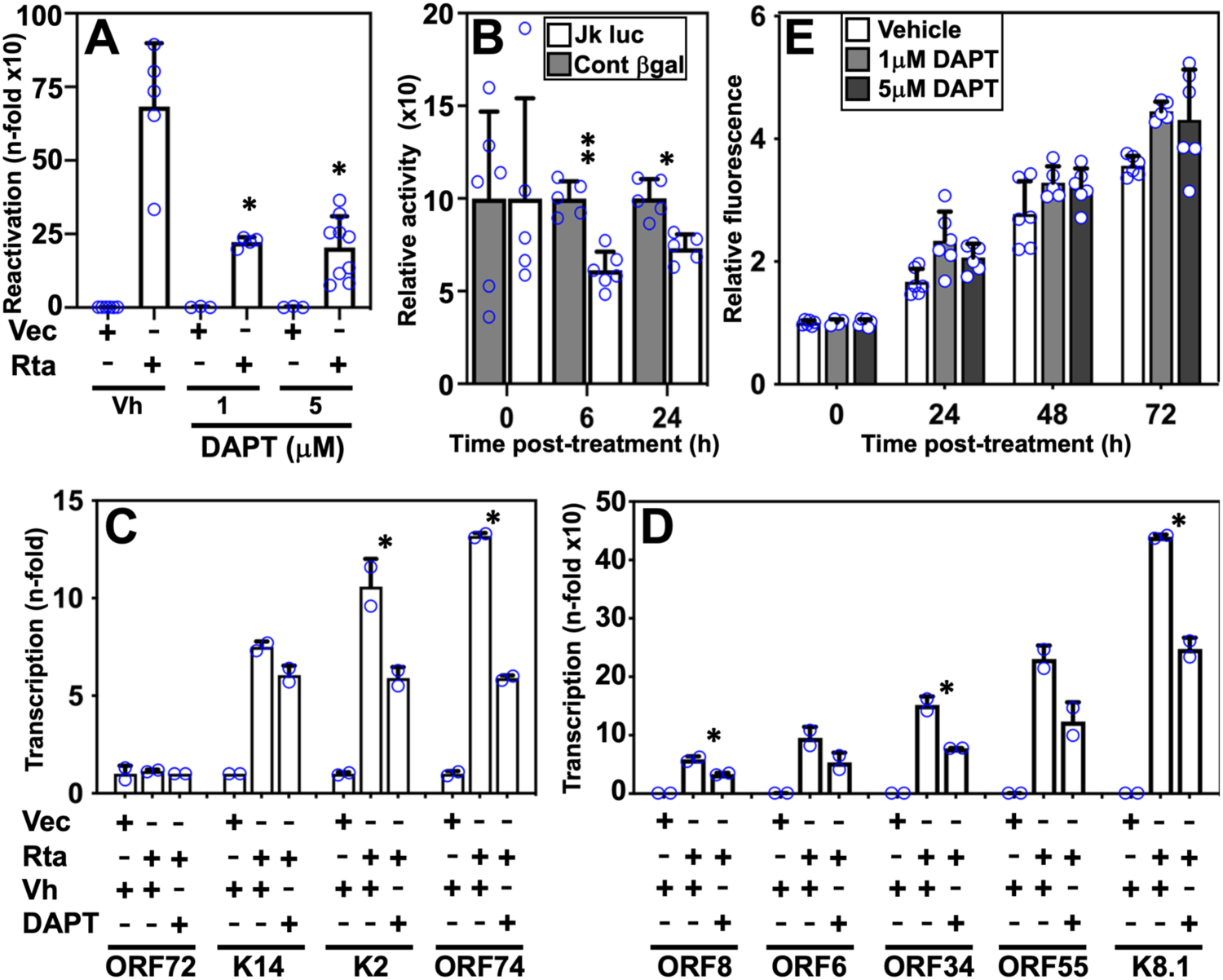
Notch inhibition reduces Rta-mediated reactivation without inhibiting cell growth. **A. Treatment with a gamma secretase inhibitor decreases viral reactivation by Rta.** Vero rKSHV.294 cells were treated with 1 or 5 uM DAPT, or vehicle control (“Vh”) 24h prior to transfection with 2.5 ug of Rta expression vector. At 72h post-transfection, infectious virus was quantitated by transferring the media to the 293 MSR tet-OFF complementing reporter cells as described in Methods. SeAP was measured in 25 uL of media per infection using the Phospha-Light System (Applied Biosystems). SeAP values from media alone were subtracted from each experimental value, and folds were calculated by normalizing to samples treated with vehicle and transfected with empty vector. *p<0.05 by t test compared to Vec + Vh. **B. Treatment with a gamma secretase inhibitor decreases activity of a Notch-dependent reporter plasmid.** Vero rKSHV.294 cells were transfected with Notch reporter plasmid pGL2-4xCSL expressing firefly luciferase (“Jk”) and pcDNA3.1-HisLacZ expressing β-galactosidase (“Cont”). Cells were treated with 5 uM DAPT or vehicle (DMSO) 24hr post-transfection, and total cellular protein was extracted. Luciferase values were normalized to βgal values at each timepoint. *p<0.0015 and **p<0.0001 by t test compared to Cont at 6h and 24h. **C. and D. Expression of KSHV transcripts are reduced by inhibiting endogenous Notch1**. Triplicate cultures of Vero rKSHV.294 cells were transfected with empty vector or Rta expression vector and treated with vehicle or DAPT (5 uM DAPT), as in Fig 2A. Total RNA was harvested 24 h after transfection and transcripts for the indicated viral genes were quantitated by real-time quantitative reverse-transcription/PCR. Transcription of the indicated viral genes was calculated by the delta-delta Ct method using cellular beta-actin transcription as the internal control for each sample and Vec + vehicle as baseline fold. One outlying value per transfection was removed using Peirce’s method. *p<0.05 by t test comparing Rta + vehicle (DMSO) to Rta + 5 uM DAPT. Each primer pair was also tested without adding reverse transcriptase to the input RNA and no amplicons were detected (D.M. Lukac, unpublished data). **E. Notch inhibition by DAPT treatment does not affect Vero rKSHV.294 cell growth.** Triplicate cultures of Vero rKSHV.294 cells were treated in two biologic replicates with vehicle or DAPT at the indicated concentrations as described in Fig 2A. At the post-treatment times indicated, cell viability was quantitated using Alamar Blue (BioRad). Mock values were averaged and subtracted from each experimental value to yield corrected values. All corrected values were then normalized to 0h values that had been set to the relative value of 1 for each treatment. The slight enhancement of cell growth rate when compared to the vehicle control was not statistically-significant.

Since reactivation is scored by transferring the supernatants from the infected Vero cells to 293 MSR tet-OFF cells, we performed a key control experiment to assess the potential effect of DAPT on the function of the 293 reporter cells. We treated virus-containing supernatant media from the experimental Vero rKSHV.294 cells with DAPT prior to incubating the supernatants with the 293 cells. There was no decrease in virus detection in the 293 cells, confirming that DAPT does not inhibit *de novo* infection by KSHV nor expression of the SEAP reporter protein (Fig S1).

To confirm that NICD activity was reduced by DAPT treatment in the infected Vero cells, we co-transfected the Vero rKSHV.294 cells with two reporter plasmids. One contained four RBP-Jk motifs arrayed in tandem upstream of a luciferase reporter gene, and the other contained the HCMV enhancer sequence lacking RBP-Jk motifs for normalization. As expected, treatment of the cells with 5 uM DAPT for 6 or 24h reduced Notch activity relative to vehicle-treated cells (Fig. 2B).

To determine if viral gene expression was similarly debilitated by inhibiting Notch activation during reactivation, we measured levels of viral transcripts in the infected Vero cells. We observed a gene specific reduction in viral RNAs (Figs 2C and D) in cells where Notch is inhibited (+DAPT) and viral reactivation is induced by ectopic Rta. The expression of K2, ORF74, ORF8, ORF6, ORF34, and K8.1 transcripts were significantly reduced by inhibiting Notch transactivation with DAPT (Figs 2C and D). ORF6 and ORF55 transcripts followed the same trend of sensitivity to Notch inhibition, but did not reach statistical significance (Fig 2D). K14 transcript was activated by Rta expression but not reduced by Notch inhibition, and ORF72 transcription was neither induced by Rta expression or repressed by DAPT pre-treatment (Fig 2C). We conclude that activated Notch cooperates with Rta to support production of infectious KSHV by transactivating specific, essential viral genes.

### Inhibiting Notch activation does not reduce growth of KSHV-infected Vero cells

Notch inhibition by DAPT can inhibit growth and induce apoptosis in KSHV-infected cells (31, 34–40). To determine the effect of DAPT on growth of the infected Vero rKSHV.294 cells, we measured cell viability by alamarBlue sensitivity. Over a 72h period of growth, cells treated with 1 uM or 5 uM DAPT maintained normal cell viability (Fig. 2E). The Fig. 2 data therefore support the conclusion that Notch inhibition by DAPT treatment specifically reduces viral reactivation without affecting Vero cell growth or viability.

### Inhibiting the Notch transactivation complex reduces KSHV reactivation

Inhibiting the gamma-secretase complex with DAPT can block the formation of several signaling substrates in addition to Notch (58). Therefore, we worked to specifically block Notch activity directly at its site of transcriptional transactivation by testing the effects of ectopically expressing dominant negative (dn) mutants of the Mastermind-like1 (MAML1) protein. Cognate MAML1 stabilizes the NICD1/RBP-Jk/DNA complex to facilitate NICD1-dependent transactivation in the nucleus (2, 3, 59, 60). Transfection of Vero rKSHV.294 cells with two different dnMAML1 expression vectors reduced Rta-mediated reactivation by 40% or 90%, respectively (Fig. 3A). As a control, we observed that dnMAML1 proteins were strongly expressed (Fig 3B).

**Fig 3.**
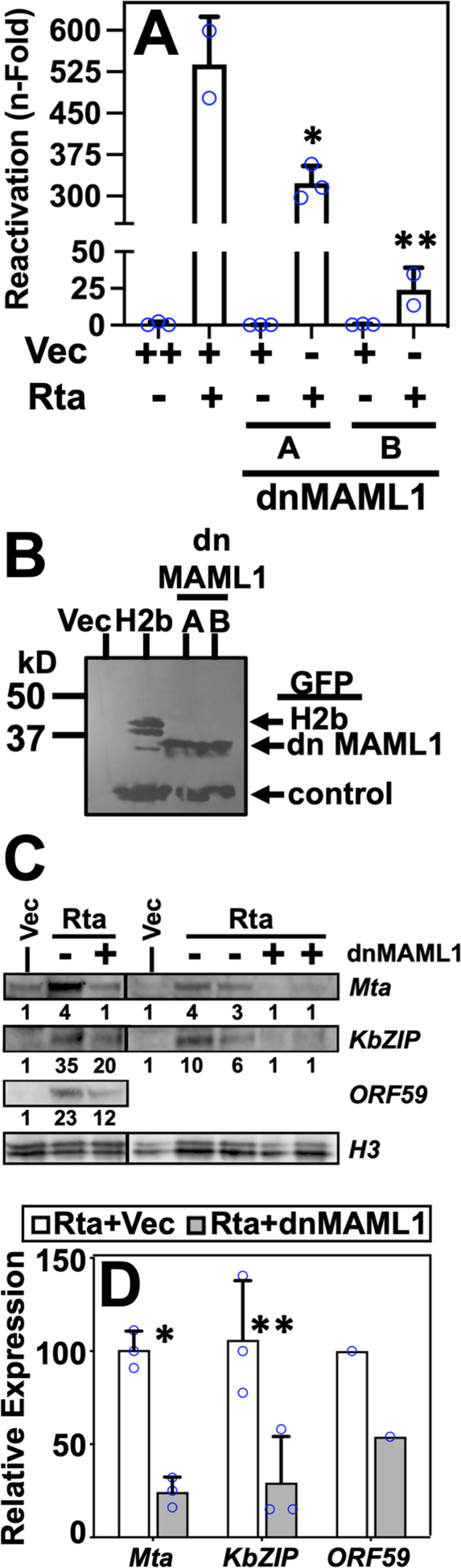
Inhibition of endogenous Notch by dominant negative MAML1 mutants inhibit viral reactivation and expression of some viral genes. **A**. Vero rKSHV.294 cells were transfected with 1ug of plasmids expressing Rta and two different dnMAML alleles (“dnMAM” “A” and “B”; each fused to GFP). Infectious virus was quantitated as in Fig. 2A. *p<0.02 and **p<0.01 by t test compared to Rta plus Vector. **B. dnMAML1 proteins are well-expressed in infected Vero rKSHV.294 cells.** Vero rKSHV.294 cells were transfected as in Fig 3A, total protein was harvested in RIPA buffer 48h later, Western blotted, and probed with GFP-specific antiserum to detect the dnMAML-GFP fusion proteins. As a control for the Western blot, plasmid expressing H2b-GFP was also transfected into cells (“H2b”). Equivalent loading of lanes was verified by probing for endogenous histone H3 (“control”). **C. Expression of KSHV proteins are reduced by inhibiting endogenous Notch1.** Expression of the KSHV proteins Mta, KbZIP, and ORF59 in extracts from cells transfected as in Fig 3B were visualized by Western blotting. Fold induction of each viral protein was determined by normalizing to expression of the host protein Histone H3 (”H3”) in each lane, then dividing by the value for Vec alone, which had been set to “1.” Those values are listed under each corresponding lane. **D. Quantitation**. The relative reduction of viral protein expression by inhibiting Notch1 was calculated by comparing each viral protein’s induction by ectopic Rta + Vec with that induced by Rta in the presence of ectopic dnMAML1 in panel B. Rta alone was set at 100. *p<0.09 and **p<0.005 by t test compared to Rta + Vec.

To determine if viral gene expression was similarly debilitated by inhibiting the Notch transactivation complex during reactivation, we measured protein levels in the infected Vero cells. We observed a gene specific reduction in viral proteins (Figs 3C and D) in cells expressing dnMAML1 relative to empty vector-transfected cells. The expression of Mta and KbZIP proteins were significantly reduced by inhibiting Notch transactivation with dnMAML1. ORF59 induction was also reduced by Notch inhibition in the single protein samples tested. Taken together with Fig 2, we conclude that activated Notch supports production of infectious KSHV by transactivating specific, essential viral genes in cooperation with Rta.

### Notch1 is necessary for optimal viral reactivation

The data in Figs. 2 and 3 show that inhibiting Notch post-translationally at two different biochemical steps of its activation dramatically reduced KSHV reactivation and support a pro-viral role for Notch signaling in viral reactivation in Vero cells. However, these experiments do not specify which of the four Notch receptors, Notch1,2,3 or 4, are required for efficient reactivation, as all four proteins are inhibited by DAPT and ectopic dnMAML1 (2, 59, 61–65).

Since Notch1 is constitutively active in Vero cells, we asked whether specific knock-down of Notch1 would affect viral reactivation. We transfected two different siRNAs to knock-down endogenous Notch1 in Vero-rKSHV.294 cells, and then induced viral reactivation by transfection of Rta expression vector (Fig. 4A) or treatment with the HDAC inhibitor valproic acid (VPA; eg. the most potent chemical inducer of viral reactivation in Vero cells (66)) (Fig. 4B). The Notch1-specific siRNAs reduced viral reactivation up to a maximum of ∼80% inhibition (“N1-B” in Fig. 4A) relative to control, scrambled siRNA (“Scr”). Knock-down of NICD1 prior to VPA treatment also reduced production of virus, but did not reach statistical significance in the small number of samples tested (Fig 4B). Knock-down of endogenous Notch1 was confirmed via Western blot (Fig 4C). These data verify that constitutively active Notch1 plays a pro-viral role in KSHV reactivation in Vero cells.

**Fig 4.**
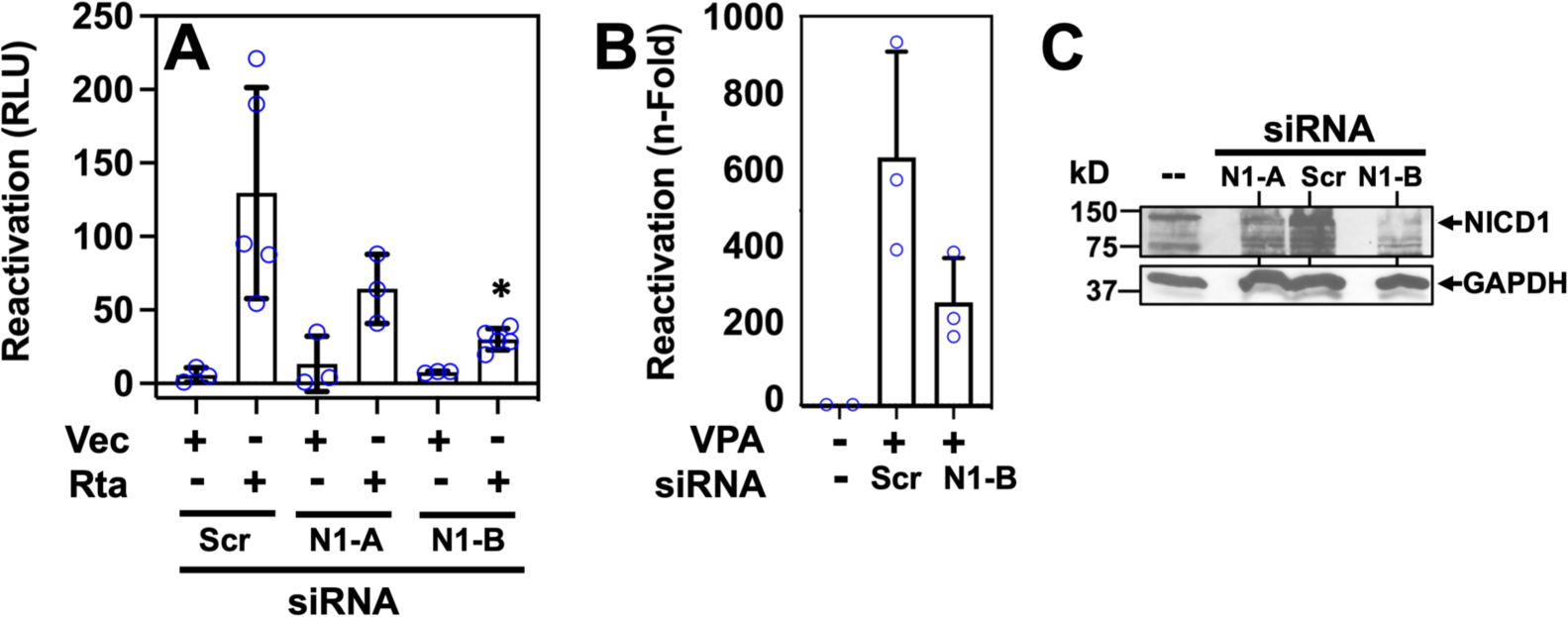
Activated Notch1 is required for optimal reactivation of KSHV. A. Reactivation by ectopic Rta. Vero rKSHV.294 cells were transfected with 30 pM of either scrambled siRNA (Scr; control), Notch 1 siRNA A (N1-A), or Notch 1 siRNA B (N1-B) in triplicate sets. 48h later, Rta expression vector or Vec (as control) was transfected into each of the siRNA-transfected cells. 48h post-transfection, virus containing media were transferred to 293 MSR tet-OFF cells. Infectious virus was quantitated as in Fig. 2A. *p<0.02 by t test compared to Rta + Scr **. B. Reactivation by VPA treatment.** Vero rKSHV.294 cells were transfected with 30pM of either scrambled siRNA (Scr; control) or Notch 1 siRNA B (N1-B). 54h later, 1 mM VPA was added to the siRNA-transfected cells. 72h post-transfection, virus containing media were transferred to 293 MSR tet-OFF cells. Infectious virus was quantitated as in Fig. 2A. **C. siRNA specifically reduces NICD1 expression.** Vero rKSHV.294 cells were transfected with 30pM of the siRNAs described in the legend to Fig. 4A. 48h post transfection, total cellular proteins were extracted in RIPA buffer, then analyzed via a 10% SDS-PAGE/western blot as described in the legend to Fig. 1A.

### Activated Notch1 associates with viral promoter DNA during KSHV reactivation

Our observations that inhibiting Notch during KSHV reactivation reduced viral gene transcription without affecting cell growth (Figs. 2 and 3) suggested that our observed proviral function of Notch1 (Fig 4) is to transactivate specific viral genes to support KSHV reactivation. Activated Notch1 (NICD1) transactivates specific cellular genes by assembling NICD1/MAML1/RBP-Jk1/DNA complexes on promoters or enhancers (2, 3, 59, 60). We and others have demonstrated that RBP-Jk1 binds to many sites on KSHV DNA (4–27), and we have mapped eighty-two RBP-Jk1 binding sites to the viral genome during reactivation (55). We selected six of these RBP-Jk1 binding sites found within 1 kb upstream of transcription start sites for KSHV genes, ie. in their presumptive promoters, and asked whether we could also detect NICD1 occupancy there using chromatin immunoprecipitation. When we induced reactivation in rKSHV.294 cells by transfecting Rta expression vector, qPCR showed that NICD1 antibody co-precipitated significantly more promoter DNA than in EV-transfected cells on the K2, ORF34, ORF6, ORF8, and the bidirectional ORF50AS/K8 promoters (Fig. 5). Transcription of four of these genes was significantly impaired when Notch activity was inhibited (Figs 2C and D). We conclude that NICD1 forms complexes with RBP-Jk and DNA to transactivate specific viral genes to promote KSHV reactivation.

**Fig. 5.**
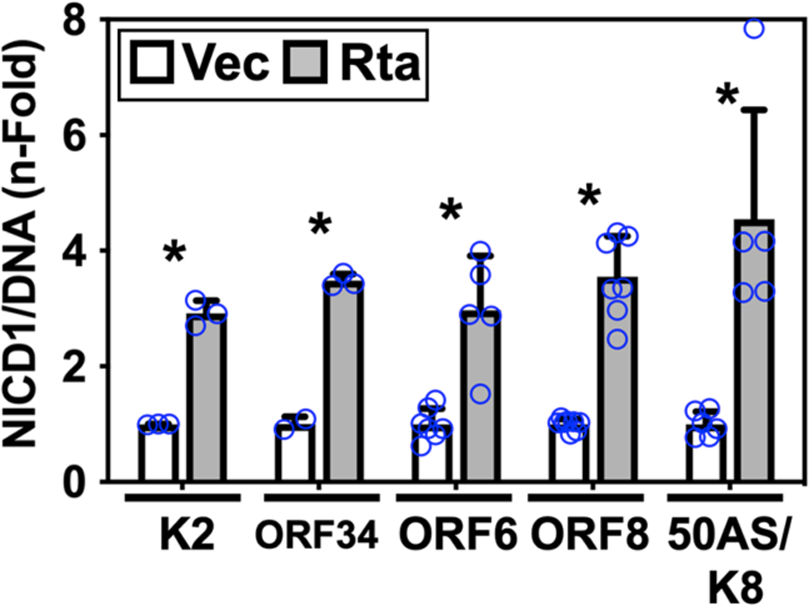
Activated Notch binds to KSHV promoters during reactivation. Triplicate cultures of Vero rKSHV.294 cells were transfected with empty vector or Rta expression vector in three experiments. Cross-linked chromatin was sonicated and immunoprecipitated (ChIP’d) with NICD1-specific antibody or non-specific IgG control. ChIP’d DNA was purified and analyzed by quantitative real-time PCR to detect sequences located within 1 kb. upstream of the transcriptional start sites of the indicated KSHV genes. Fold increases in ChIP’d promoters were calculated by the delta-delta Ct method using non-specific IgG as the internal control for each sample and EV transfection as baseline fold. Folds were normalized by comparison to NICD1 expression in transfected samples. *p<0.05 by t test comparing Rta to Vec for each primer pair.

### Notch is pro-viral in PEL cells

Notch1 is constitutively active in PEL cells (36–38). To determine whether Notch1 also supports reactivation in these cells, we reactivated virus with VPA in the presence of 16, 32, or 40 uM of DAPT and quantitated vDNA accumulation and protein expression. DAPT reduced intracellular vDNA accumulation up to a maximum of ∼80% relative to VPA plus vehicle control (“Vh”) (Fig 6A). DAPT also reduced the percentage of K8.1-positive cells (Fig. 6B); as K8.1 is a true-late gene, this result was consistent with the effect of DAPT on vDNA accumulation.

**Fig. 6.**
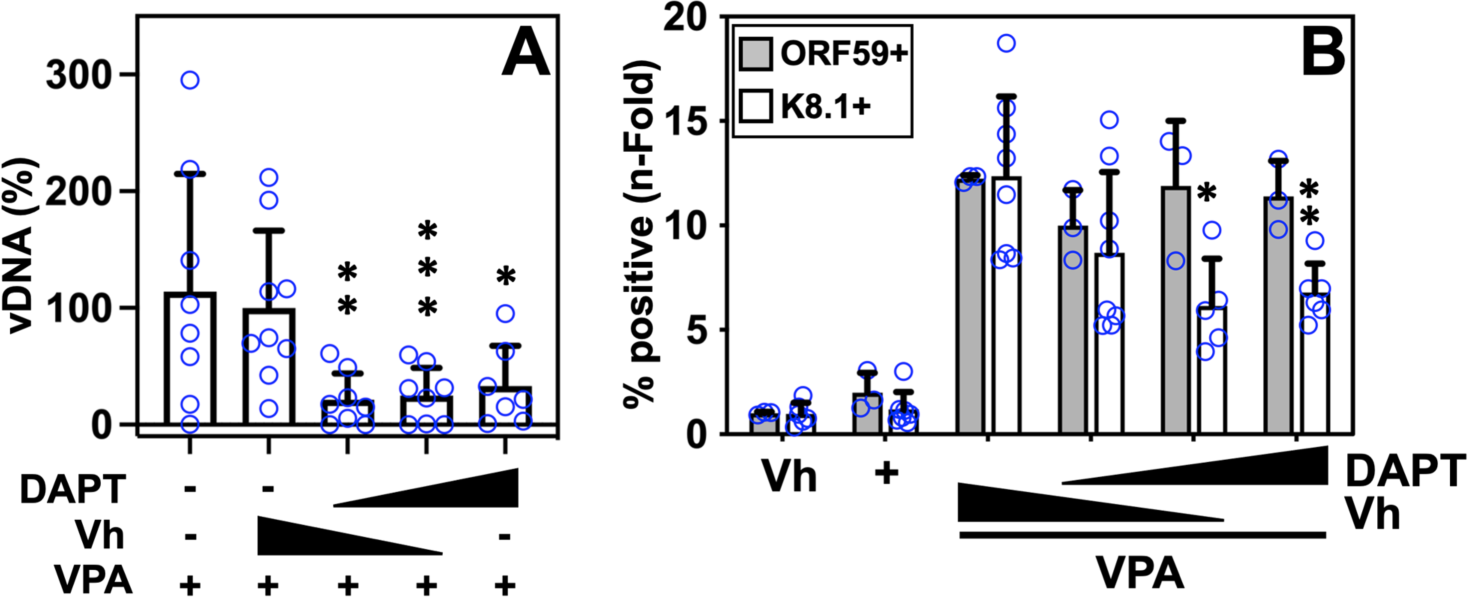
Activated Notch1 is required for optimal reactivation of KSHV in infected PEL cells. A. DAPT inhibits vDNA accumulation in PEL cells. Triplicate cultures of KSHV-infected BC3 cells were treated with vehicle alone (“Vh”) or containing 16, 32, or 40 uM DAPT for 65h. Total DNA was purified using the FlexiGene DNA kit (Qiagen). Intracellular viral DNA was measured by real-time quantitative (q) PCR using SYBR Green PCR Master Mix (Thermo Scientific) and GAPDH was the internal control. Fold-DNA was calculated using the ΔCt method and all values were normalized to that of VPA + vehicle alone, which was set to 100%. *p<0.03, **p<0.009, and ***p<0.006 by t test compared to VPA+Veh. Results from three biologic replicates were combined. **B. Specific expression of KSHV genes is reduced by Notch inhibition in PEL cells.** Triplicate cultures of KSHV-infected BCBL1 cells were treated with 1 mM VPA and vehicle (“Vh”) or DAPT (16, 32, or 40 uM). 48 or 72h post-treatment, ORF59 and K8.1 were detected by indirect immunofluorescence, respectively. The percentages of ORF59 or K8.1 positive cells were quantitated, and fold-percent positive cells were calculated by normalizing to samples treated with vehicle alone. Results from three biologic replicates were combined. *p<0.007 and **p<0.005 by t test compared to VPA+Veh.

Interestingly, we did not observe an effect of DAPT treatment on the percentage of ORF59-positive cells, perhaps because this immunofluorescence assay in PEL cells fails to detect modest effects on the ORF59 amounts or signal intensity (Fig 6B). In Vero cells, western blotting suggested that Notch inhibition reduced overall ORF59 protein expression (Fig 3D).

In an SLK cell model of KSHV infection, Notch was shown to inhibit reactivation and reduce expression of productive cycle genes, including Rta (33). In contrast, we did not observe a statistically significant effect of DAPT treatment on Rta expression in any of the experiments we performed in Vero and PEL cells (J. Decotiis-Mauro and D.M. Lukac, unpublished data). Therefore, based on the DE kinetics of expression of many of the Notch-sensitive viral genes (Figs 2, 3, 5, and 6), we conclude that Notch cooperates with Rta to promote reactivation at an early time point that precedes vDNA replication.

### Ectopic, activated Notch isoforms 1 to 4 are insufficient to reactivate KSHV from latency

In our experiments using the infected Vero cell system, as well as experiments in PEL cells, we demonstrated that ectopic NICD1 is incapable of productively reactivating KSHV, despite widespread RBP-Jk binding to the latent viral genome (6, 9, 30). Mammals express four Notch receptors; although other Notch isoforms are commonly constitutively-expressed in KSHV-infected cells (31, 34, 36–38) it is possible that their endogenous levels are insufficient to drive viral reactivation.

Notch receptors 1 to 4 encode all of the functional domains present in NICD1, except for Notch3 and 4, which lack the Notch transactivation domain (reviewed in (2, 67, 68)). Importantly, all four NICDs contain the RBP-Jk-interaction domain and activate promoters containing RBP-Jk DNA motifs. Target selectivity of each NICD differs based on promoter contexts of the RBP-Jk motifs and the cellular contexts in which each receptor is activated (2, 48, 68–80). The sequences and arrangement of RBP-Jk motifs in individual promoters determine the magnitude of transactivation by particular NICD proteins. Moreover, since each Notch receptor can cross-talk with other signaling pathways and transcription factor networks, the co-occupancy of promoters with particular NICD/RBP-Jk complexes and heterologous transcription factors can also determine the responsiveness of a gene to a specific Notch receptor and its respective NICD.

To test whether ectopic Notch isoforms 2, 3, and 4 could reactivate KSHV, we transfected vectors that express NICD 2, 3, and 4 into Vero rKSHV.294 cells. As expected, ectopic Rta, the positive control, gave a 500-fold increase in KSHV reactivation as measured by SeAP activity (Fig. 7A). However, little to no virus was detectable from cells transfected with any of the NICD expression vectors, similar to effects in cells transfected with the Rta transcriptionally deficient mutant ORF50ΔSTAD (1.7-fold; Fig. 7A). All four isoforms were thus insufficient to reactivate KSHV from Vero cells, despite their robust expression levels (Fig 7B).

**Fig. 7.**
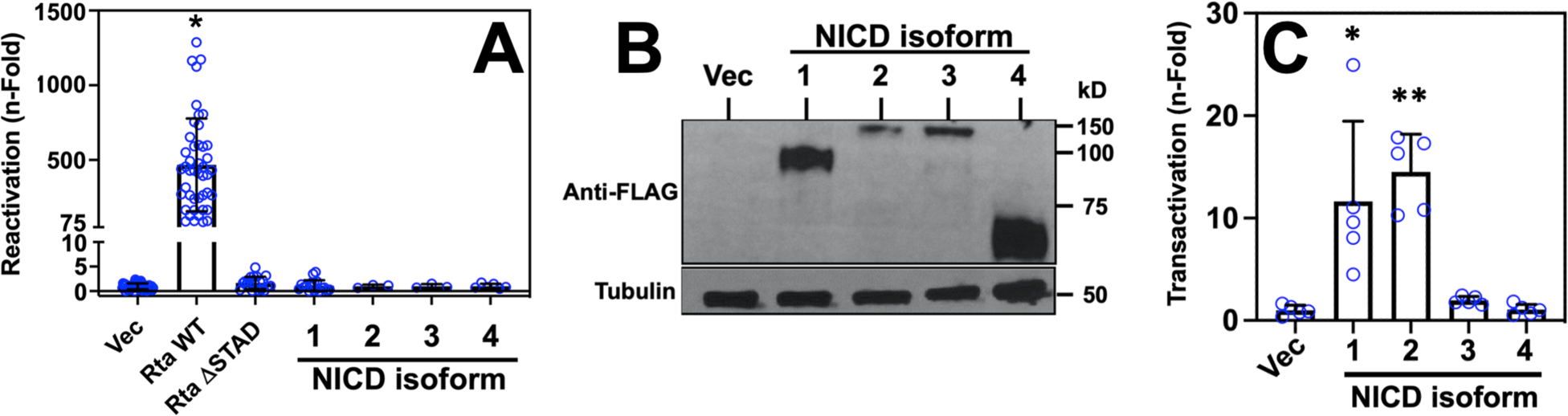
Ectopic NICD 1-4 are not sufficient to induce KSHV reactivation. **A**. Vero rKSHV.294 cells were transfected with 2.5 ug plasmid DNA expressing each of the indicated proteins and incubated for 72h at which time virus containing media were transferred to 293 MSR tet-OFF cells. Infectious virus was quantitated as in the legend to Fig. 2A, using the Great EscAPe SeAP kit (Takara). *p<0.0002 by t test compared to Vec. Data points shown are from one to ten biologic replicates containing two to three technical replicates for each experimental condition. **B. All four constitutively active Notch isoforms are expressed ectopically in transfected Vero rKSHV.294 cells.** Cells were transfected as in the legend to Fig 7A, and total proteins were harvested in RIPA buffer 48h later, Western blotted, and probed with FLAG-tag and tubulin-specific antibodies. All Notch isoforms were observed at the appropriate apparent MWs (48). **C. Ectopic Notch isoforms 1 and 2 are transactivation competent in Vero rKSHV.294 cells.** Triplicate cultures of Vero rKSHV.294 cells were transfected with 0.25 ug of the Notch reporter plasmid 4xCBS expressing firefly luciferase, control plasmid pcDNA3.1-HisLacZ expressing β-galactosidase (βgal) and 3 ug of each of the four Notch isoform vectors. Extracts were made 48 h post-transfection, and raw luciferase values were corrected to β-galactosidase. Folds were calculated by normalization to samples transfected with luciferase reporter and empty vector (Vec) alone. *p<0.02 and **p<0.0001 by t test compared to Vec.

### Ectopic NICD1 and 2 are transactivation competent in KSHV-infected Vero cells

It is possible that the activated Notch proteins are unable to reactivate KSHV because they are not transcriptionally active in Vero rKSHV.294 cells. To assess their activities, we co-transfected Vero rKSHV.294 cells with a reporter vector containing four tandem RBP-Jκ sites, and each of the four NICD expression vectors at the same amounts as in Fig 7A. We observed significant transactivation of this promoter by both NICD1 and 2, but not NICD3 or 4 (Fig 7C). Taken together with Fig. 7A, these data confirm that NICD1 and 2 are incapable of reactivating KSHV despite being transcriptionally active. However, the inability of NICD3 or 4 to activate transcription in these cells provides a potential explanation for their inability to reactivate KSHV from latency.

### NICD mutants of varying activity are insufficient to reactivate KSHV from latency

NICD1 dosage has dramatic effects on biologic phenotypes. Since endogenous, constitutively-active Notch1 is abundant in the infected Vero cells (Fig 1), it was possible that ectopic NICD1 failed to reactivate KSHV (Fig 7A) due to excessive Notch activity. To vary the gain-of-function activity of Notch1 in the infected Vero cells, we expressed a set of four Notch1 alleles known to have varying intrinsic signaling and transactivating potencies. Notch1-P12 and Notch1-L1594P, identified in clinical T cell acute lymphoblastic leukemia (T-ALL) lymphomas, are full-length Notch1 receptors expressed from the pCMV vector that each produce constitutively-active NICD1 due to mutations in the Notch1 heterodimerization domain that promote cleavage of the mutant proteins (49, 81)(Fig S2A). Both Notch1 mutants are weaker transcriptional transactivators than WT pcDNA3-NICD1, which expresses NICD1 lacking the transmembrane domain and protease cleavage sites and bypasses the requirement for proteolysis to generate the active NICD1 transactivator (49, 81). The fourth clone, pCMV-NICD1-ΔPEST, expresses a mutant of NICD1 protein that also lacks the PEST proteasomal degradation domain, conferring increased stability to NICD1 and stronger, more sustained transactivation. We transfected vectors expressing each of these alleles into the Vero rKSHV.294 cells and found that none could reactivate the virus, similar to the result with cognate NICD1 (“WT”; Fig S2B). These data further verify that NICD1 is insufficient to reactivate KSHV from latency, regardless of its inherent potency or expression level.

### NICD1 cooperates with VPA in reactivation

Our data showed that endogenous Notch1 is constitutively active in infected Vero cells and supports optimal viral reactivation that is induced by VPA treatment or by ectopic Rta expression. However, increasing activated Notch1 amounts alone, via ectopic NICD1 expression, was insufficient to productively reactivate KSHV (Figs 7 and S2). To determine whether ectopic NICD1 could cooperate with VPA treatment, we transfected the Vero rKSHV.294 cells with the NICD1 expression vector alone or with 1 mM VPA treatment. In positive control samples treated with VPA alone, we observe approximately 350-fold reactivation (Fig 8). As expected, ectopic expression of NICD1 alone was insufficient to produce virus (Fig 8). However, expression of ectopic NICD1 in VPA-treated cells significantly enhanced reactivation to about 700-fold (Fig 8), supporting a cooperative role for NICD1 and VPA. The NICD1 mutants, NICD1ΔPEST, P12, and L1594P, also increased VPA-stimulated reactivation, but did not reach statistical significance due to the small sample size (Fig S2C).

**Fig. 8.**
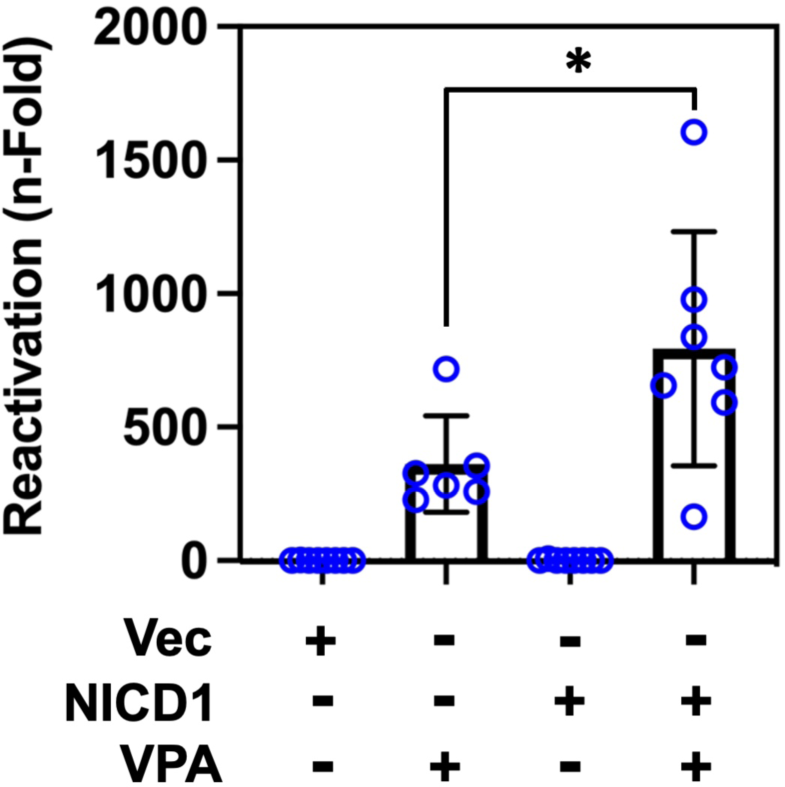
Ectopic NICD1 cooperates with VPA in KSHV reactivation. Vero rKSHV.294 cells were transfected with 1 ug plasmid DNA expressing NICD1 (“N1”) or empty vector. 18 h post-transfection, cells were treated with 1 mM VPA. 72 h post-treatment, virus containing media were transferred to 293 MSR tet-OFF cells. Infectious virus was quantitated as in Fig 2A. *p<0.05 by t test.

## Discussion

In this work, we demonstrate that ectopic expression of activated Notch1 is insufficient to productively reactivate KSHV from latency (Fig 7). NICD1’s reactivation insufficiency confirms our earlier results from ectopic expression of NICD1 in PEL and Vero cells (6, 30), and agrees with similar results in which NICD1 was conditionally-induced ectopically in a stably-transfected PEL cell line (29). To ensure that this negative result is not due to an inadequate dose of ectopic NICD1, we show that three additional Notch1 alleles that encode constitutively active NICD1 proteins with varied intrinsic transcriptional potency also fail to reactivate KSHV from latency (Fig S2). Our work is the first to test all four of the NICD isoforms in this capacity, and conclusively establishes that ectopic NICD2 is also not capable of reactivating the virus. While ectopic expression of NICD isoforms 3 and 4 also did not result in production of infectious virus, we are unable to conclude that NICD3 and 4 are similarly insufficient to reactivate KSHV since we could not determine that they are transcriptionally active in our experiments (Fig 7C). However, since the Notch isoforms have been shown to have different promoter transactivation specificities (48, 82–84), it is possible that NICD3 and 4 are, in fact, active in Vero cells, but are simply incapable of activating the 4x tandem RBP-Jk motif (4x CBS) reporter plasmid. NICD1 and 2 have been suggested to be functionally interchangeable in other biologic systems (85–87).

Despite Notch’s inability to reactivate latent KSHV alone, we show that constitutive Notch1 activity is required for robust reactivation of the virus. All three of our strategies that inhibited endogenous Notch activity in the Vero cells decreased Rta or VPA-induced reactivation over a range of 60-90% (Figs 2 to 4 and 6). Two of the strategies, DAPT treatment and dnMAML1 expression, are well-established strategies to broadly inhibit all four Notch isoforms (2, 59, 61–65). DAPT blocks gamma-secretase activation of endogenous Notch and dnMAML1 interferes with the formation of Notch transactivation complexes (46, 88–90). However, neither strategy is strictly Notch-specific, because each also interferes with other cellular processes that require gamma-secretase activity or cognate MAML1 (58, 60). To inhibit Notch1 specifically, we knocked-down its expression using two different transfected siRNAs (Fig. 4). Inhibiting Notch with dnMAML1 reduced reactivation more dramatically than did siRNA knock-down, suggesting that endogenous Notch 2, 3, and 4 may also have pro-viral functions (compare Figs 3 and 4). Our Notch inhibition studies focused primarily on quantitating production of infectious virus from latently infected Vero cells, but we also observed that inhibiting Notch in PEL cells reduced synthesis of intracellular viral DNA (Fig 6A). Those data suggest that Notch activity in both cell lines have similar effects on viral infection. We note a single contradictory study that concluded that ectopic NICD1 reactivated KSHV (32).

It is well-established that Notch signaling is critical for cell and animal models of KSHV pathogenesis; however, our data are the first to demonstrate that Notch activation is necessary for robust production of infectious virus. Productive cycle gene expression is also seen in the context of sustained Notch activation that is required for the endothelial to mesenchymal transition of KSHV-infected cells (31, 35, 39, 40). Growth and survival of KSHV-infected cells also depend on Notch activity: proliferation rates of KSHV-infected B cells were proportional to Notch expression levels, and Notch inhibition reduced cellular cyclin D1 expression and arrested cells in G1 (37). Moreover, pharmacologic Notch inhibition induced apoptosis of infected SLK, B, and primary KS tumor cells (31, 34, 35, 37–40) and induced necrosis of PEL xenografts in NOD/SCID mice (36). Although Notch1 expression is insufficient to productively reactivate KSHV, ectopic Notch1 could induce expression of a subset of productive cycle viral genes in PEL cells (29). Importantly, the infected Vero cell model used in our work is ideal for uncoupling the effect of Notch on cell growth from its effect on viral reactivation; in our work, we used DAPT concentrations that significantly reduced production of infectious virus without affecting host cell doubling time or survival (Fig 2E). This pro-viral role of Notch follows the paradigm established in foundational tumor virus studies in which cellular oncogenic pathways manipulated by SV40 or Adenovirus also support productive viral infections (91).

Our data raised two intriguing questions: how does activated Notch contribute to reactivation and why is it not sufficient to reactivate KSHV alone? Since complete Notch ablation is inconsistent with survival of KSHV-infected cells, one apparent contribution of activated Notch is to maintain survival of cells in which KSHV is reactivating (31, 34, 36–40). To this end, we note that none of our strategies to reduce Notch activity were 100% effective in debilitating virus production. However, our observation that DAPT treatment and partial NICD1 knock-down can debilitate reactivation without reducing cell growth suggests a function of Notch that is complementary to Rta and independent of cell survival.

Our data also show that NICD1 is transactivation competent in latently-infected Vero cells (Fig 7C), as has also been demonstrated in latently-infected PEL cells (29). Thus, all of the components sufficient to form the Notch1 transactivation complex must be expressed and functional in KSHV-infected cells (reviewed in (2, 3, 92)). Therefore, we believe that Notch’s insufficiency to reactivate KSHV derives from a limit to gene-specific transactivation that is relieved when Rta is expressed. Importantly, one critical gene which ectopic NICD1 fails to activate in Vero and PEL cells is Rta (J. DeCotiis-Mauro and D.M. Lukac, unpublished data (29)). Therefore, when endogenous Rta is induced by VPA, or Rta is expressed ectopically, endogenous NICD1 then likely transactivates specific viral genes during reactivation. Indeed, Notch forms complexes with vDNA in the presumptive promoters of five viral genes (Fig 5), and Notch inhibition is associated with reduction in expression of four of the genes and others not tested (Figs 2 and 3). Importantly, of the NICD1 targets identified herein, ORF8/gB, ORF34, K8.1, ORF57/Mta, K8/KbZIP, and ORF59 are all essential for viral production in at least one cell type (93–100). Our observation that broad Notch inhibitors have greater anti-viral potency than specific Notch1 inhibition (compare Figs 2A and 3A to Fig 4A) suggests that at least one additional Notch isoform also supports optimal KSHV reactivation.

We interpret our data as revealing that Notch activity had no influence on productive reactivation unless Rta was expressed. Activated Notch has no intrinsic DNA binding ability, but instead relies upon the cellular DNA binding protein RBP-Jk to specify NICD1’s target genes (2, 3, 101). One function of Rta which can profoundly influence the ability of NICD1 to transactivate specific genes is Rta’s ability to stimulate RBP-Jk DNA binding (6, 9, 28). Our lab modeled this mechanism by showing that NICD1 can transactivate the Mta promoter only if a truncated form of Rta, lacking its transactivation domain, stimulates RBP-Jk binding to the promoter (6). Studying the intact virus, we showed that KSHV reactivation is associated with dramatic re-localization of RBP-Jk to unique sites on the virus genome (28). The extent of dynamic RBP-Jk binding to the virus can be contrasted with that observed on the cell genome. In uninfected cells, RBP-Jk binding sites on the cellular genome are classified as static or dynamic, and NICD1 can stimulate RBP-Jk DNA binding to cellular genomes (102–105). However, on the KSHV genome, the proportion of dynamic RBP-Jk sites in our studies vastly exceed the proportion of dynamic sites reported on cellular genomes (28, 102, 103, 106–109). In fact, the number of reactivation-specific RBP-Jk sites on the virus, and the number of RBP-Jk DNA motifs on the virus, exceed the number of KSHV ORFs (28). Moreover, Rta is not found bound to the virus at every RBP-Jk site (28). We hypothesize that NICD1/MAML1/RBP-Jk complexes transactivate specific essential viral promoters and cooperate with Rta to support KSHV reactivation (Fig 8). During latency, in the absence of Rta expression, NICD1/RBP-Jk fails to form a competent transactivation complex on at least one essential viral promoter. Unlike NICD1, Rta also binds DNA independent of RBP-Jk for transactivation and replication (110–113); these mechanistic differences probably also contribute to Notch’s insufficiency in reactivation.

Based on experimental data, we have published two mechanisms that describe how Rta regulates RBP-Jk DNA binding: (1) Rta directly binds to RBP-Jk, and (2) Rta induces expression of additional cellular proteins during reactivation which in turn bind to DNA adjacent to RBP-Jk (6, 9, 27, 28). The latter mechanism is consistent with our work and others that demonstrated that *de novo* gene expression downstream of Rta is required for robust reactivation (5, 27). Thus, NICD1 transactivation in KSHV-infected cells is determined by the fate of the infection (latent vs. reactivating) and intracellular context, as well as the DNA context of individual RBP-Jk binding sites. Similar context-dependent determinants have been invoked to explain cell-specific variations in NICD1 transactivation and phenotypic consequences in uninfected cells (48, 73–78, 109, 114–117). The influence of cellular context on Notch function includes interactions between the four Notch isoforms, including competition, cooperation, or redundancy (83, 85, 86, 118–131).

In Notch cancer biology, cell context is a major determinant of the striking observation that Notch can be either tumor suppressive or oncogenic in different malignancies (reviewed in (2, 80, 132, 133)). The ideal DNA context for an NICD1-responsive RBP-Jk site is a head-to-head configuration with a second RBP-Jk site spaced 15-17 nts apart (69–72); remarkably, only six pairs of RBP-Jk motifs in the KSHV genome are found in that DNA context (J. DeCotiis-Mauro and D.M. Lukac, unpublished data). It is likely that the mechanisms underlying contextual variations in Notch signaling outcomes in uninfected cells are functionally similar in KSHV infection, but uniquely regulated by KSHV proteins like Rta. Moreover, cell context may be responsible for the discordance of our observations with a publication that showed that NICD1 inhibits KSHV reactivation in SLK cells (33).

In summary, herein we demonstrate the significance of constitutive Notch1 activity in the cellular context of productive KSHV reactivation. The essentiality of Rta/RBP-Jk/DNA complexes in driving reactivation is well-established, but this current study demonstrates that Notch contributes to reactivation by forming promoter-specific NICD1/RBP-Jk/DNA complexes, also. The KSHV genome contains >100 RBP-Jk motifs of varying sequence found in varying DNA contexts that we expect to respond differently to Rta and/or NICD1 to determine the success of a reactivation signal leading to virus production. Moreover, our data indicate that Notch isoforms other than the well-studied Notch1 may help regulate viral infection. This diversity in the molecular players underlies the consequences of RBP-Jk-dependent gene expression in KSHV infection and host cell pathology. Our data are consistent with the idea that the Notch-dependance of viral reactivation is a complex network that, when revealed, will provide important new insights into Notch-dependent signaling in mammalian cells.

## Acknowledgements

We gratefully thank Drs. Tom Kadesch, Jeff Vieira, Ethel Cesarman, Signe V. Horn Pederson, Geoff Wahl, Rhett Kovall, Raphael Kopan, Jon C. Aster, and Spyros Artavanis-Tsakonas for cells, antibody, and plasmids. The work was supported by NIH Grants AI117127 and AI150230.

## References

1. Watanabe T, Sugimoto A, Hosokawa K, Fujimuro M. 2018. Signal Transduction Pathways Associated with KSHV-Related Tumors. Adv Exp Med Biol 1045:321–355.

2. Aster JC, Pear WS, Blacklow SC. 2017. The Varied Roles of Notch in Cancer. Annu Rev Pathol 12:245–275.

3. Giaimo BD, Gagliani EK, Kovall RA, Borggrefe T. 2021. Transcription Factor RBPJ as a Molecular Switch in Regulating the Notch Response, p 9–30. *In* Reichrath J, Reichrath S (ed), Notch Signaling in Embryology and Cancer: Notch Signaling in Cancer doi:10.1007/978-3-030-55031-8_2. Springer International Publishing, Cham.

4. Campbell M, Watanabe T, Nakano K, Davis RR, Lyu Y, Tepper CG, Durbin-Johnson B, Fujimuro M, Izumiya Y. 2018. KSHV episomes reveal dynamic chromatin loop formation with domain-specific gene regulation. Nat Commun 9:49.

5. Papp B, Motlagh N, Smindak RJ, Jin Jang S, Sharma A, Alonso JD, Toth Z. 2019. Genome-Wide Identification of Direct RTA Targets Reveals Key Host Factors for Kaposi’s Sarcoma-Associated Herpesvirus Lytic Reactivation. J Virol 93.

6. Carroll KD, Bu W, Palmeri D, Spadavecchia S, Lynch SJ, Marras SA, Tyagi S, Lukac DM. 2006. Kaposi’s Sarcoma-associated herpesvirus lytic switch protein stimulates DNA binding of RBP-Jk/CSL to activate the Notch pathway. J Virol 80:9697–709.

7. Liang Y, Chang J, Lynch S, Lukac DM, Ganem D. 2002. The lytic switch protein of KSHV activates gene expression via functional interaction with RBP-Jk, the target of the Notch signaling pathway. Genes Dev 16:1977–89.

8. Liang Y, Ganem D. 2003. Lytic but not latent infection by Kaposi’s sarcoma-associated herpesvirus requires host CSL protein, the mediator of Notch signaling. Proc Natl Acad Sci U S A 100:8490–5.

9. Palmeri D, Carroll KD, Gonzalez-Lopez O, Lukac DM. 2011. Kaposi’s sarcoma-associated herpesvirus Rta tetramers make high-affinity interactions with repetitive DNA elements in the Mta promoter to stimulate DNA binding of RBP-Jk/CSL. J Virol 85:11901–15.

10. Hilton IB, Dittmer DP. 2012. Quantitative analysis of the bidirectional viral G-protein-coupled receptor and lytic latency-associated nuclear antigen promoter of Kaposi’s sarcoma-associated herpesvirus. J Virol 86:9683–95.

11. Bu W, Carroll KD, Palmeri D, Lukac DM. 2007. The Kaposi’s Sarcoma-associated Herpesvirus/Human herpesvirus-8 ORF50/Rta Lytic Switch Protein Functions as a Tetramer. J Virol 81:5788–5806.

12. Lukac D, Garibyan L, Kirshner J, Palmeri D, Ganem D. 2001. DNA binding by the Kaposi’s sarcoma-associated herpesvirus lytic switch protein is necessary for transcriptional activation of two viral delayed early promoters. J Virol 75:6786–99.

13. Lukac DM, Kirshner JR, Ganem D. 1999. Transcriptional activation by the product of open reading frame 50 of Kaposi’s sarcoma-associated herpesvirus is required for lytic viral reactivation in B cells. J Virol 73:9348–61.

14. Lukac DM, Renne R, Kirshner JR, Ganem D. 1998. Reactivation of Kaposi’s sarcoma-associated herpesvirus infection from latency by expression of the ORF 50 transactivator, a homolog of the EBV R protein. Virology 252:304–312.

15. Wang Y, Yuan Y. 2006. Essential role of RBP-Jkappa in activation of the K8 delayed-early promoter of Kaposi’s sarcoma-associated herpesvirus by ORF50/RTA. Virology.

16. Persson LM, Wilson AC. 2010. Wide-scale use of Notch signaling factor CSL/RBP-Jkappa in RTA-mediated activation of Kaposi’s sarcoma-associated herpesvirus lytic genes. J Virol 84:1334–47.

17. Scholz BA, Harth-Hertle ML, Malterer G, Haas J, Ellwart J, Schulz TF, Kempkes B. 2013. Abortive lytic reactivation of KSHV in CBF1/CSL deficient human B cell lines. PLoS Pathog 9:e1003336.

18. Lan K, Kuppers DA, Verma SC, Robertson ES. 2004. Kaposi’s sarcoma-associated herpesvirus-encoded latency-associated nuclear antigen inhibits lytic replication by targeting Rta: a potential mechanism for virus-mediated control of latency. J Virol 78:6585–94.

19. Liu Y, Cao Y, Liang D, Gao Y, Xia T, Robertson ES, Lan K. 2008. Kaposi’s sarcoma-associated herpesvirus RTA activates the processivity factor ORF59 through interaction with RBP-Jkappa and a cis-acting RTA responsive element. Virology 380:264–75.

20. Liang Y, Ganem D. 2004. RBP-J (CSL) is essential for activation of the K14/vGPCR promoter of Kaposi’s sarcoma-associated herpesvirus by the lytic switch protein RTA. J Virol 78:6818–26.

21. Chang PJ, Boonsiri J, Wang SS, Chen LY, Miller G. 2010. Binding of RBP-Jkappa (CSL) protein to the promoter of the Kaposi’s sarcoma-associated herpesvirus ORF47 (gL) gene is a critical but not sufficient determinant of transactivation by ORF50 protein. Virology 398:38–48.

22. Izumiya Y, Izumiya C, Hsia D, Ellison TJ, Luciw PA, Kung HJ. 2009. NF-kappaB serves as a cellular sensor of Kaposi’s sarcoma-associated herpesvirus latency and negatively regulates K-Rta by antagonizing the RBP-Jkappa coactivator. J Virol 83:4435–46.

23. Sun R, LIn SF, Gradoville L, Yuan Y, Zhu F, Miller G. 1998. A viral gene that activates lytic cycle expression of Kaposi’s sarcoma-associated herpesvirus. Proc Natl Acad Sci USA 95:10866–10871.

24. Gradoville L, Gerlach J, Grogan E, Shedd D, Nikiforow S, Metroka C, Miller G. 2000. Kaposi’s sarcoma-associated herpesvirus open reading frame 50/Rta protein activates the entire lytic cycle in the HH-B2 primary effusion lymphoma cell line. J Virol 74:6207–6212.

25. Nakamura H, Lu M, Gwack Y, Souvlis J, Zeichner SL, Jung JU. 2003. Global changes in Kaposi’s sarcoma-associated virus gene expression patterns following expression of a tetracycline-inducible Rta transactivator. J Virol 77:4205–20.

26. Xu Y, AuCoin DP, Huete AR, Cei SA, Hanson LJ, Pari GS. 2005. A Kaposi’s sarcoma-associated herpesvirus/human herpesvirus 8 ORF50 deletion mutant is defective for reactivation of latent virus and DNA replication. J Virol 79:3479–87.

27. Bu W, Palmeri D, Krishnan R, Marin R, Aris VM, Soteropoulous P, Lukac DM. 2008. Identification of Direct Transcriptional Targets of the KSHV Rta Lytic Switch Protein by conditional nuclear localization. J Virol 82:10709–10723.

28. Gonzalez-Lopez O DJ, Goyeneche C, Mello H, Vicente-Ortiz BA, Shin HJ, Driscoll KE, Du P, Palmeri D, Lukac DM. 2019. A herpesvirus transactivator and cellular POU proteins extensively regulate DNA binding of the host Notch signaling protein RBP-Jκ to the virus genome. J Biol Chem 294:13073–13092.

29. Chang H, Dittmer DP, Chul SY, Hong Y, Jung JU. 2005. Role of Notch signal transduction in Kaposi’s sarcoma-associated herpesvirus gene expression. J Virol 79:14371–82.

30. DeCotiis JL, Ortiz NC, Vega BA, Lukac DM. 2017. An easily transfectable cell line that produces an infectious reporter virus for routine and robust quantitation of Kaposi’s sarcoma-associated herpesvirus reactivation. J Virol Methods 247:99–106.

31. Cheng F, Pekkonen P, Laurinavicius S, Sugiyama N, Henderson S, Gunther T, Rantanen V, Kaivanto E, Aavikko M, Sarek G, Hautaniemi S, Biberfeld P, Aaltonen L, Grundhoff A, Boshoff C, Alitalo K, Lehti K, Ojala PM. 2011. KSHV-initiated notch activation leads to membrane-type-1 matrix metalloproteinase-dependent lymphatic endothelial-to-mesenchymal transition. Cell Host Microbe 10:577–90.

32. Lan K, Murakami M, Choudhuri T, Kuppers DA, Robertson ES. 2006. Intracellular-activated Notch1 can reactivate Kaposi’s sarcoma-associated herpesvirus from latency. Virology 351:393–403.

33. Li S, Hu H, He Z, Liang D, Sun R, Lan K. 2016. Fine-Tuning of the Kaposi’s Sarcoma-Associated Herpesvirus Life Cycle in Neighboring Cells through the RTA-JAG1-Notch Pathway. PLOS Pathogens 12:e1005900.

34. Curry CL, Reed LL, Golde TE, Miele L, Nickoloff BJ, Foreman KE. 2005. Gamma secretase inhibitor blocks Notch activation and induces apoptosis in Kaposi’s sarcoma tumor cells. Oncogene 24:6333–44.

35. Emuss V, Lagos D, Pizzey A, Gratrix F, Henderson SR, Boshoff C. 2009. KSHV manipulates Notch signaling by DLL4 and JAG1 to alter cell cycle genes in lymphatic endothelia. PLoS Pathog 5:e1000616.

36. Lan K, Murakami M, Bajaj B, Kaul R, He Z, Gan R, Feldman M, Robertson ES. 2009. Inhibition of KSHV-infected primary effusion lymphomas in NOD/SCID mice by gamma-secretase inhibitor. Cancer Biol Ther 8:2136–43.

37. Lan K, Choudhuri T, Murakami M, Kuppers DA, Robertson ES. 2006. Intracellular activated Notch1 is critical for proliferation of Kaposi’s sarcoma-associated herpesvirus-associated B-lymphoma cell lines in vitro. J Virol 80:6411–9.

38. Lan K, Verma SC, Murakami M, Bajaj B, Kaul R, Robertson ES. 2007. Kaposi’s sarcoma herpesvirus-encoded latency-associated nuclear antigen stabilizes intracellular activated Notch by targeting the Sel10 protein. Proc Natl Acad Sci U S A 104:16287–92.

39. Liu R, Li X, Tulpule A, Zhou Y, Scehnet JS, Zhang S, Lee JS, Chaudhary PM, Jung J, Gill PS. 2010. KSHV-induced notch components render endothelial and mural cell characteristics and cell survival. Blood 115:887–95.

40. Gasperini P, Espigol-Frigole G, McCormick PJ, Salvucci O, Maric D, Uldrick TS, Polizzotto MN, Yarchoan R, Tosato G. 2012. Kaposi sarcoma herpesvirus promotes endothelial-to-mesenchymal transition through Notch-dependent signaling. Cancer Res 72:1157–69.

41. Gould F, Harrison SM, Hewitt EW, Whitehouse A. 2009. Kaposi’s Sarcoma-Associated Herpesvirus RTA Promotes Degradation of the Hey1 Repressor Protein through the Ubiquitin Proteasome Pathway. Journal of Virology 83:6727–6738.

42. Green MR, Sambrook J, Sambrook J. 2012. Molecular cloning: a laboratory manual, 4th ed. Cold Spring Harbor Laboratory Press, Cold Spring Harbor, N.Y.

43. Lukac DM, Renne R, Kirshner JR, Ganem D. 1998. Reactivation of Kaposi’s Sarcoma-Associated Herpesvirus Infection from Latency by Expression of the ORF 50 Transactivator, a Homolog of the EBV R Protein. Virology 252:304–312.

44. Horn S, Kobberup S, Jørgensen MC, Kalisz M, Klein T, Kageyama R, Gegg M, Lickert H, Lindner J, Magnuson MA, Kong Y-Y, Serup P, Ahnfelt-Rønne J, Jensen JN. 2012. <em>Mind bomb 1</em> is required for pancreatic β-cell formation. Proceedings of the National Academy of Sciences 109:7356–7361.

45. Kanda T, Sullivan KF, Wahl GM. 1998. Histone-GFP fusion protein enables sensitive analysis of chromosome dynamics in living mammalian cells. Curr Biol 8:377–85.

46. Saxena MT, Schroeter EH, Mumm JS, Kopan R. 2001. Murine Notch Homologs (N1&#x2013;4) Undergo Presenilin-dependent Proteolysis *. Journal of Biological Chemistry 276:40268–40273.

47. Yuan Z, Friedmann DR, VanderWielen BD, Collins KJ, Kovall RA. 2012. Characterization of CSL (CBF-1, Su(H), Lag-1) mutants reveals differences in signaling mediated by Notch1 and Notch2. J Biol Chem 287:34904–16.

48. Ong CT, Cheng HT, Chang LW, Ohtsuka T, Kageyama R, Stormo GD, Kopan R. 2006. Target selectivity of vertebrate notch proteins. Collaboration between discrete domains and CSL-binding site architecture determines activation probability. J Biol Chem 281:5106–19.

49. Weng AP, Ferrando AA, Lee W, Morris JPt, Silverman LB, Sanchez-Irizarry C, Blacklow SC, Look AT, Aster JC. 2004. Activating mutations of NOTCH1 in human T cell acute lymphoblastic leukemia. Science 306:269–71.

50. Gantt S, Carlsson J, Ikoma M, Gachelet E, Gray M, Geballe AP, Corey L, Casper C, Lagunoff M, Vieira J. 2011. The HIV protease inhibitor nelfinavir inhibits Kaposi’s sarcoma-associated herpesvirus replication in vitro. Antimicrobial agents and chemotherapy 55:2696–703.

51. Carroll KD, Khadim F, Spadavecchia S, Palmeri D, Lukac DM. 2007. Direct interactions of Kaposi’s sarcoma-associated herpesvirus/human herpesvirus 8 ORF50/Rta protein with the cellular protein octamer-1 and DNA are critical for specifying transactivation of a delayed-early promoter and stimulating viral reactivation. J Virol 81:8451–67.

52. Kirshner JR, Lukac DM, Chang J, Ganem D. 2000. Kaposi’s Sarcoma-Associated Herpesvirus Open Reading Frame 57 Encodes a Posttranscriptional Regulator with Multiple Distinct Activities. J Virol 74:3586–3597.

53. Bruce AG, Barcy S, DiMaio T, Gan E, Garrigues HJ, Lagunoff M, Rose TM. 2017. Quantitative Analysis of the KSHV Transcriptome Following Primary Infection of Blood and Lymphatic Endothelial Cells. Pathogens 6.

54. Journo G, Tushinsky C, Shterngas A, Avital N, Eran Y, Karpuj MV, Frenkel-Morgenstern M, Shamay M. 2018. Modulation of Cellular CpG DNA Methylation by Kaposi’s Sarcoma-Associated Herpesvirus. J Virol 92.

55. Gonzalez-Lopez O, Goyeneche, C, DeCotiis, J Vicente-Ortiz, BA, Shin H, Driscoll, K, Du P, Palmeri, D, and DM Lukac. 2019. Viral Rta and cellular POU proteins regulate dynamic, genome-wide DNA binding of the Notch pathway protein Recombination Signal Binding Protein (RBP)-Jk during reactivation of Kaposi’s sarcoma-associated herpesvirus (KSHV). Submitted.

56. Nelson JD, Denisenko O, Bomsztyk K. 2006. Protocol for the fast chromatin immunoprecipitation (ChIP) method. Nature Protocols 1:179–185.

57. Ross S. 2003. Peirce’s criterion for the elimination of suspect experimental data. J Eng Technol 20.

58. Güner G, Lichtenthaler SF. 2020. The substrate repertoire of γ-secretase/presenilin. Seminars in Cell & Developmental Biology 105:27–42.

59. Weng AP, Nam Y, Wolfe MS, Pear WS, Griffin JD, Blacklow SC, Aster JC. 2003. Growth Suppression of Pre-T Acute Lymphoblastic Leukemia Cells by Inhibition of Notch Signaling. Molecular and Cellular Biology 23:655–664.

60. Zema S, Pelullo M, Nardozza F, Felli MP, Screpanti I, Bellavia D. 2020. A Dynamic Role of Mastermind-Like 1: A Journey Through the Main (Path)ways Between Development and Cancer. Frontiers in Cell and Developmental Biology 8.

61. Maillard I, Weng AP, Carpenter AC, Rodriguez CG, Sai H, Xu L, Allman D, Aster JC, Pear WS. 2004. Mastermind critically regulates Notch-mediated lymphoid cell fate decisions. Blood 104:1696–1702.

62. Saravanamuthu SS, Gao CY, Zelenka PS. 2009. Notch signaling is required for lateral induction of Jagged1 during FGF-induced lens fiber differentiation. Dev Biol 332:166–76.

63. Bagheri L, Pellati A, Rizzo P, Aquila G, Massari L, De Mattei M, Ongaro A. 2018. Notch pathway is active during osteogenic differentiation of human bone marrow mesenchymal stem cells induced by pulsed electromagnetic fields. J Tissue Eng Regen Med 12:304–315.

64. Lilly B, Kennard S. 2009. Differential gene expression in a coculture model of angiogenesis reveals modulation of select pathways and a role for Notch signaling. Physiol Genomics 36:69–78.

65. Sastre M, Steiner H, Fuchs K, Capell A, Multhaup G, Condron MM, Teplow DB, Haass C. 2001. Presenilin-dependent γ-secretase processing of β-amyloid precursor protein at a site corresponding to the S3 cleavage of Notch. EMBO reports 2:835–841.

66. Shin HJ, Decotiis J, Giron M, Palmeri D, Lukac DM. 2014. Histone Deacetylase Classes I and II Regulate Kaposi’s Sarcoma-Associated Herpesvirus Reactivation. J Virol 88:1281–92.

67. You W-K, Schuetz TJ, Lee SH. 2023. Targeting the DLL/Notch Signaling Pathway in Cancer: Challenges and Advances in Clinical Development. Molecular Cancer Therapeutics 22:3–11.

68. Hori K, Sen A, Artavanis-Tsakonas S. 2013. Notch signaling at a glance. J Cell Sci 126:2135–40.

69. Nam Y, Sliz P, Pear WS, Aster JC, Blacklow SC. 2007. Cooperative assembly of higher-order Notch complexes functions as a switch to induce transcription. Proc Natl Acad Sci U S A 104:2103–8.

70. Nam Y, Sliz P, Song L, Aster JC, Blacklow SC. 2006. Structural basis for cooperativity in recruitment of MAML coactivators to Notch transcription complexes. Cell 124:973–83.

71. Arnett KL, Hass M, McArthur DG, Ilagan MX, Aster JC, Kopan R, Blacklow SC. 2010. Structural and mechanistic insights into cooperative assembly of dimeric Notch transcription complexes. Nat Struct Mol Biol 17:1312–7.

72. Liu H, Chi AW, Arnett KL, Chiang MY, Xu L, Shestova O, Wang H, Li YM, Bhandoola A, Aster JC, Blacklow SC, Pear WS. 2010. Notch dimerization is required for leukemogenesis and T-cell development. Genes Dev 24:2395–407.

73. Castro B, Barolo S, Bailey AM, Posakony JW. 2005. Lateral inhibition in proneural clusters: cis-regulatory logic and default repression by Suppressor of Hairless. Development 132:3333–3344.

74. Nellesen DT, Lai EC, Posakony JW. 1999. Discrete enhancer elements mediate selective responsiveness of enhancer of split complex genes to common transcriptional activators. Dev Biol 213:33–53.

75. Wang H, Zou J, Zhao B, Johannsen E, Ashworth T, Wong H, Pear WS, Schug J, Blacklow SC, Arnett KL, Bernstein BE, Kieff E, Aster JC. 2011. Genome-wide analysis reveals conserved and divergent features of Notch1/RBPJ binding in human and murine T-lymphoblastic leukemia cells. Proc Natl Acad Sci U S A 108:14908–13.

76. Swanson CI, Evans NC, Barolo S. 2010. Structural rules and complex regulatory circuitry constrain expression of a Notch– and EGFR-regulated eye enhancer. Dev Cell 18:359–70.

77. Cave JW, Loh F, Surpris JW, Xia L, Caudy MA. 2005. A DNA transcription code for cell-specific gene activation by notch signaling. Curr Biol 15:94–104.

78. Basak O, Giachino C, Fiorini E, MacDonald HR, Taylor V. 2012. Neurogenic Subventricular Zone Stem/Progenitor Cells Are Notch1-Dependent in Their Active But Not Quiescent State. The Journal of Neuroscience 32:5654–5666.

79. Bray SJ. 2016. Notch signalling in context. Nature Reviews Molecular Cell Biology 17:722–735.

80. Arruga F, Vaisitti T, Deaglio S. 2018. The NOTCH Pathway and Its Mutations in Mature B Cell Malignancies. Frontiers in Oncology 8.

81. Chiang MY, Xu L, Shestova O, Histen G, L’Heureux S, Romany C, Childs ME, Gimotty PA, Aster JC, Pear WS. 2008. Leukemia-associated NOTCH1 alleles are weak tumor initiators but accelerate K-ras-initiated leukemia. J Clin Invest 118:3181–94.

82. Beatus P, Lundkvist J, Oberg C, Pedersen K, Lendahl U. 2001. The origin of the ankyrin repeat region in Notch intracellular domains is critical for regulation of HES promoter activity. Mech Dev 104:3–20.

83. James AC, Szot JO, Iyer K, Major JA, Pursglove SE, Chapman G, Dunwoodie SL. 2014. Notch4 reveals a novel mechanism regulating Notch signal transduction. Biochim Biophys Acta 1843:1272–84.

84. Yamaguchi N, Oyama T, Ito E, Satoh H, Azuma S, Hayashi M, Shimizu K, Honma R, Yanagisawa Y, Nishikawa A, Kawamura M, Imai J-i, Ohwada S, Tatsuta K, Inoue J-i, Semba K, Watanabe S. 2008. NOTCH3 Signaling Pathway Plays Crucial Roles in the Proliferation of ErbB2-Negative Human Breast Cancer Cells. Cancer Research 68:1881–1888.

85. Liu Z, Brunskill E, Varnum-Finney B, Zhang C, Zhang A, Jay PY, Bernstein I, Morimoto M, Kopan R. 2015. The intracellular domains of Notch1 and Notch2 are functionally equivalent during development and carcinogenesis. Development 142:2452–2463.

86. Giachino C, Boulay JL, Ivanek R, Alvarado A, Tostado C, Lugert S, Tchorz J, Coban M, Mariani L, Bettler B, Lathia J, Frank S, Pfister S, Kool M, Taylor V. 2015. A Tumor Suppressor Function for Notch Signaling in Forebrain Tumor Subtypes. Cancer Cell 28:730–742.

87. Rossi D, Trifonov V, Fangazio M, Bruscaggin A, Rasi S, Spina V, Monti S, Vaisitti T, Arruga F, Famà R, Ciardullo C, Greco M, Cresta S, Piranda D, Holmes A, Fabbri G, Messina M, Rinaldi A, Wang J, Agostinelli C, Piccaluga PP, Lucioni M, Tabbò F, Serra R, Franceschetti S, Deambrogi C, Daniele G, Gattei V, Marasca R, Facchetti F, Arcaini L, Inghirami G, Bertoni F, Pileri SA, Deaglio S, Foà R, Dalla-Favera R, Pasqualucci L, Rabadan R, Gaidano G. 2012. The coding genome of splenic marginal zone lymphoma: activation of NOTCH2 and other pathways regulating marginal zone development. Journal of Experimental Medicine 209:1537–1551.

88. De Strooper B, Annaert W, Cupers P, Saftig P, Craessaerts K, Mumm JS, Schroeter EH, Schrijvers V, Wolfe MS, Ray WJ, Goate A, Kopan R. 1999. A presenilin-1-dependent γ-secretase-like protease mediates release of Notch intracellular domain. Nature 398:518–522.

89. Lin SE, Oyama T, Nagase T, Harigaya K, Kitagawa M. 2002. Identification of new human mastermind proteins defines a family that consists of positive regulators for notch signaling. J Biol Chem 277:50612–20.

90. Wu L, Sun T, Kobayashi K, Gao P, Griffin JD. 2002. Identification of a family of mastermind-like transcriptional coactivators for mammalian notch receptors. Mol Cell Biol 22:7688–700.

91. DiMaio D. 2019. Small size, big impact: how studies of small DNA tumour viruses revolutionized biology. Philos Trans R Soc Lond B Biol Sci 374:20180300.

92. Siebel C, Lendahl U. 2017. Notch Signaling in Development, Tissue Homeostasis, and Disease. Physiol Rev 97:1235–1294.

93. Zhou Y, Tian X, Wang S, Gao M, Zhang C, Ma J, Cheng X, Bai L, Qin HB, Luo MH, Qin Q, Jiang B, Lan K, Zhang J. 2024. Palmitoylation of KSHV pORF55 is required for Golgi localization and efficient progeny virion production. PLoS Pathog 20:e1012141.

94. Dollery SJ, Santiago-Crespo RJ, Chatterjee D, Berger EA. 2019. Glycoprotein K8.1A of Kaposi’s Sarcoma-Associated Herpesvirus Is a Critical B Cell Tropism Determinant Independent of Its Heparan Sulfate Binding Activity. J Virol 93.

95. Han Z, Swaminathan S. 2006. Kaposi’s sarcoma-associated herpesvirus lytic gene ORF57 is essential for infectious virion production. J Virol 80:5251–60.

96. Majerciak V, Pripuzova N, McCoy JP, Gao SJ, Zheng ZM. 2007. Targeted disruption of Kaposi’s sarcoma-associated herpesvirus ORF57 in the viral genome is detrimental for the expression of ORF59, K8alpha, and K8.1 and the production of infectious virus. J Virol 81:1062–71.

97. Wang Y, Sathish N, Hollow C, Yuan Y. 2011. Functional characterization of Kaposi’s sarcoma-associated herpesvirus open reading frame K8 by bacterial artificial chromosome-based mutagenesis. J Virol 85:1943–57.

98. Krishnan HH, Sharma-Walia N, Zeng L, Gao SJ, Chandran B. 2005. Envelope glycoprotein gB of Kaposi’s sarcoma-associated herpesvirus is essential for egress from infected cells. J Virol 79:10952–67.

99. Nishimura M, Watanabe T, Yagi S, Yamanaka T, Fujimuro M. 2017. Kaposi’s sarcoma-associated herpesvirus ORF34 is essential for late gene expression and virus production. Sci Rep 7:329.

100. Peng C, Chen J, Tang W, Liu C, Chen X. 2014. Kaposi’s sarcoma-associated herpesvirus ORF6 gene is essential in viral lytic replication. PLoS One 9:e99542.

101. Sprinzak D, Blacklow SC. 2021. Biophysics of Notch Signaling. Annu Rev Biophys 50:157–189.

102. Wang H, Zang C, Taing L, Arnett KL, Wong YJ, Pear WS, Blacklow SC, Liu XS, Aster JC. 2014. NOTCH1-RBPJ complexes drive target gene expression through dynamic interactions with superenhancers. Proc Natl Acad Sci U S A 111:705–10.

103. Castel D, Mourikis P, Bartels SJ, Brinkman AB, Tajbakhsh S, Stunnenberg HG. 2013. Dynamic binding of RBPJ is determined by Notch signaling status. Genes Dev 27:1059–71.

104. Krejci A, Bray S. 2007. Notch activation stimulates transient and selective binding of Su(H)/CSL to target enhancers. Genes Dev 21:1322–7.

105. Gomez-Lamarca MJ, Falo-Sanjuan J, Stojnic R, Abdul Rehman S, Muresan L, Jones ML, Pillidge Z, Cerda-Moya G, Yuan Z, Baloul S, Valenti P, Bystricky K, Payre F, O’Holleran K, Kovall R, Bray SJ. 2018. Activation of the Notch Signaling Pathway In Vivo Elicits Changes in CSL Nuclear Dynamics. Developmental Cell 44:611–623.e7.

106. Zhao B, Zou J, Wang H, Johannsen E, Peng CW, Quackenbush J, Mar JC, Morton CC, Freedman ML, Blacklow SC, Aster JC, Bernstein BE, Kieff E. 2011. Epstein-Barr virus exploits intrinsic B-lymphocyte transcription programs to achieve immortal cell growth. Proc Natl Acad Sci U S A 108:14902–7.

107. Severson E, Arnett KL, Wang H, Zang C, Taing L, Liu H, Pear WS, Shirley Liu X, Blacklow SC, Aster JC. 2017. Genome-wide identification and characterization of Notch transcription complex– binding sequence-paired sites in leukemia cells. Science Signaling 10:eaag1598.

108. Ryan RJH, Petrovic J, Rausch DM, Zhou Y, Lareau CA, Kluk MJ, Christie AL, Lee WY, Tarjan DR, Guo B, Donohue LKH, Gillespie SM, Nardi V, Hochberg EP, Blacklow SC, Weinstock DM, Faryabi RB, Bernstein BE, Aster JC, Pear WS. 2017. A B Cell Regulome Links Notch to Downstream Oncogenic Pathways in Small B Cell Lymphomas. Cell Reports 21:784–797.

109. Yashiro-Ohtani Y, Wang H, Zang C, Arnett KL, Bailis W, Ho Y, Knoechel B, Lanauze C, Louis L, Forsyth KS, Chen S, Chung Y, Schug J, Blobel GA, Liebhaber SA, Bernstein BE, Blacklow SC, Liu XS, Aster JC, Pear WS. 2014. Long-range enhancer activity determines <em>Myc</em> sensitivity to Notch inhibitors in T cell leukemia. Proceedings of the National Academy of Sciences doi:10.1073/pnas.1407079111:201407079.

110. Song M, Brown H, Wu T-T, Sun R. 2001. Transcription activation of polyadenylated nuclear RNA by Rta in Human herpesvirus 8/Kaposi’s Sarcoma-associated herpesvirus. J Virol 75:3129–40.

111. Chen J, Ye F, Xie J, Kuhne K, Gao SJ. 2009. Genome-wide identification of binding sites for Kaposi’s sarcoma-associated herpesvirus lytic switch protein, RTA. Virology 386:290–302.

112. Wang Y, Li H, Chan MY, Zhu FX, Lukac DM, Yuan Y. 2004. Kaposi’s sarcoma-associated herpesvirus ori-Lyt-dependent DNA replication: cis-acting requirements for replication and ori-Lyt-associated RNA transcription. J Virol 78:8615–29.

113. Rossetto CC, Susilarini NK, Pari GS. 2011. Interaction of Kaposi’s sarcoma-associated herpesvirus ORF59 with oriLyt is dependent on binding with K-Rta. J Virol 85:3833–41.

114. Cooper MT, Tyler DM, Furriols M, Chalkiadaki A, Delidakis C, Bray S. 2000. Spatially restricted factors cooperate with notch in the regulation of Enhancer of split genes. Dev Biol 221:390–403.

115. Takebayashi K, Sasai Y, Sakai Y, Watanabe T, Nakanishi S, Kageyama R. 1994. Structure, chromosomal locus, and promoter analysis of the gene encoding the mouse helix-loop-helix factor HES-1. Negative autoregulation through the multiple N box elements. J Biol Chem 269:5150–6.

116. Stoeck A, Lejnine S, Truong A, Pan L, Wang H, Zang C, Yuan J, Ware C, MacLean J, Garrett-Engele PW, Kluk M, Laskey J, Haines BB, Moskaluk C, Zawel L, Fawell S, Gilliland G, Zhang T, Kremer BE, Knoechel B, Bernstein BE, Pear WS, Liu XS, Aster JC, Sathyanarayanan S. 2014. Discovery of biomarkers predictive of GSI response in triple-negative breast cancer and adenoid cystic carcinoma. Cancer Discov 4:1154–67.

117. Krejcí A, Bernard F, Housden BE, Collins S, Bray SJ. 2009. Direct response to Notch activation: signaling crosstalk and incoherent logic. Sci Signal 2:ra1.

118. Choi SH, Severson E, Pear WS, Liu XS, Aster JC, Blacklow SC. 2017. The common oncogenomic program of NOTCH1 and NOTCH3 signaling in T-cell acute lymphoblastic leukemia. PLOS ONE 12:e0185762.

119. Bellavia D, Campese AF, Checquolo S, Balestri A, Biondi A, Cazzaniga G, Lendahl U, Fehling HJ, Hayday AC, Frati L, von Boehmer H, Gulino A, Screpanti I. 2002. Combined expression of pTα and Notch3 in T cell leukemia identifies the requirement of preTCR for leukemogenesis. Proceedings of the National Academy of Sciences 99:3788–3793.

120. Engler A, Rolando C, Giachino C, Saotome I, Erni A, Brien C, Zhang R, Zimber-Strobl U, Radtke F, Artavanis-Tsakonas S, Louvi A, Taylor V. 2018. Notch2 Signaling Maintains NSC Quiescence in the Murine Ventricular-Subventricular Zone. Cell Reports 22:992–1002.

121. Meester JAN, Verstraeten A, Alaerts M, Schepers D, Van Laer L, Loeys BL. 2019. Overlapping but distinct roles for NOTCH receptors in human cardiovascular disease. Clinical Genetics 95:85–94.

122. Beatus P, Lundkvist J, Oberg C, Lendahl U. 1999. The notch 3 intracellular domain represses notch 1-mediated activation through Hairy/Enhancer of split (HES) promoters. Development 126:3925–3935.

123. Beà S, Valdés-Mas R, Navarro A, Salaverria I, Martín-Garcia D, Jares P, Giné E, Pinyol M, Royo C, Nadeu F, Conde L, Juan M, Clot G, Vizán P, Di Croce L, Puente DA, López-Guerra M, Moros A, Roue G, Aymerich M, Villamor N, Colomo L, Martínez A, Valera A, Martín-Subero JI, Amador V, Hernández L, Rozman M, Enjuanes A, Forcada P, Muntañola A, Hartmann EM, Calasanz MJ, Rosenwald A, Ott G, Hernández-Rivas JM, Klapper W, Siebert R, Wiestner A, Wilson WH, Colomer D, López-Guillermo A, López-Otín C, Puente XS, Campo E. 2013. Landscape of somatic mutations and clonal evolution in mantle cell lymphoma. Proceedings of the National Academy of Sciences 110:18250–18255.

124. Ables JL, DeCarolis NA, Johnson MA, Rivera PD, Gao Z, Cooper DC, Radtke F, Hsieh J, Eisch AJ. 2010. Notch1 Is Required for Maintenance of the Reservoir of Adult Hippocampal Stem Cells. The Journal of Neuroscience 30:10484–10492.

125. Nwabo Kamdje AH, Bassi G, Pacelli L, Malpeli G, Amati E, Nichele I, Pizzolo G, Krampera M. 2012. Role of stromal cell-mediated Notch signaling in CLL resistance to chemotherapy. Blood Cancer Journal 2:e73–e73.

126. Demehri S, Turkoz A, Manivasagam S, Yockey Laura J, Turkoz M, Kopan R. 2012. Elevated Epidermal Thymic Stromal Lymphopoietin Levels Establish an Antitumor Environment in the Skin. Cancer Cell 22:494–505.

127. Boucher JM, Harrington A, Rostama B, Lindner V, Liaw L. 2013. A Receptor-Specific Function for Notch2 in Mediating Vascular Smooth Muscle Cell Growth Arrest Through Cyclin-dependent Kinase Inhibitor 1B. Circulation Research 113:975–985.

128. Southgate L, Sukalo M, Karountzos ASV, Taylor EJ, Collinson CS, Ruddy D, Snape KM, Dallapiccola B, Tolmie JL, Joss S, Brancati F, Digilio MC, Graul-Neumann LM, Salviati L, Coerdt W, Jacquemin E, Wuyts W, Zenker M, Machado RD, Trembath RC. 2015. Haploinsufficiency of the NOTCH1 Receptor as a Cause of Adams&#x2013;Oliver Syndrome With Variable Cardiac Anomalies. Circulation: Cardiovascular Genetics 8:572–581.

129. Kawai H, Kawaguchi D, Kuebrich BD, Kitamoto T, Yamaguchi M, Gotoh Y, Furutachi S. 2017. Area-Specific Regulation of Quiescent Neural Stem Cells by Notch3 in the Adult Mouse Subependymal Zone. The Journal of Neuroscience 37:11867–11880.

130. Arasada RR, Amann JM, Rahman MA, Huppert SS, Carbone DP. 2014. EGFR blockade enriches for lung cancer stem-like cells through Notch3-dependent signaling. Cancer Research doi:10.1158/0008-5472.Can-13-3724:canres.3724.2013.

131. Kofler NM, Cuervo H, Uh MK, Murtomäki A, Kitajewski J. 2015. Combined deficiency of Notch1 and Notch3 causes pericyte dysfunction, models CADASIL and results in arteriovenous malformations. Scientific Reports 5:16449.

132. Parmigiani E, Taylor V, Giachino C. 2020. Oncogenic and Tumor-Suppressive Functions of NOTCH Signaling in Glioma. Cells 9:2304.

133. D’Assoro AB, Leon-Ferre R, Braune EB, Lendahl U. 2022. Roles of Notch Signaling in the Tumor Microenvironment. Int J Mol Sci 23.

134. Schneider CA, Rasband WS, Eliceiri KW. 2012. NIH Image to ImageJ: 25 years of image analysis. Nature Methods 9:671–675.

