## Supplemental Figure Legends for "The cellular Notch1 Protein Promotes KSHV reactivation in an Rta-dependent manner"

**Figure S1. Treatment with DAPT does not affect KSHV infectivity.** Viral infected media from Vero rKSHV.294 cells transfected with 2.5 ug of genomic Rta were untreated or incubated with DMSO or 5 uM DAPT for 1h at 37°C prior to transferring to 293 MSR tet-OFF cells. Infection was quantitated as described in the Fig 2A legend.

**Fig. S2. Ectopic NICD1 alleles with varying intrinsic activities are not sufficient to induce KSHV reactivation.** **A.** Vero rKSHV.294 cells were transfected with 2.5 ug plasmid DNA expressing each of the indicated proteins. Total proteins from duplicate cultures were harvested in RIPA buffer. Equal amounts of protein from each sample, as determined by Bradford Assay, were separated by SDS-PAGE, Western blotted, and probed with Notch1-specific antibody (Santa Cruz). kD=MW markers. **B.** Vero rKSHV.294 cells were transfected with 2.5 ug plasmid DNA expressing each of the indicated proteins and incubated for 72h at which time virus containing media were transferred to 293 MSR tet-OFF cells. Infectious virus was quantitated as in Fig. 2A, using the Great EscAPe SeAP kit (Takara). *p<0.0006 and **p<0.0002 by t test compared to Empty vector (“Vec”). **C.** Vero rKSHV.294 cells were transfected with 1 ug plasmid DNA expressing each of the indicated NICD1 alleles or empty vector. 18 h post-transfection, cells were treated with 1 mM VPA. 72 h post-treatment, virus containing media were transferred to 293 MSR tet-OFF cells. Infectious virus was quantitated as in Fig 2A. Relative light units (RLU) from VPA-treated cells expressing each NICD1 allele were divided by RLU from VPA-treated cells transfected with empty vector and plotted. *p<0.05 compared to Vec+VPA by t test.
