## Supplementary figures and images for "The cellular Notch1 Protein Promotes KSHV reactivation in an Rta-dependent manner"

### Supplemental Figure S1

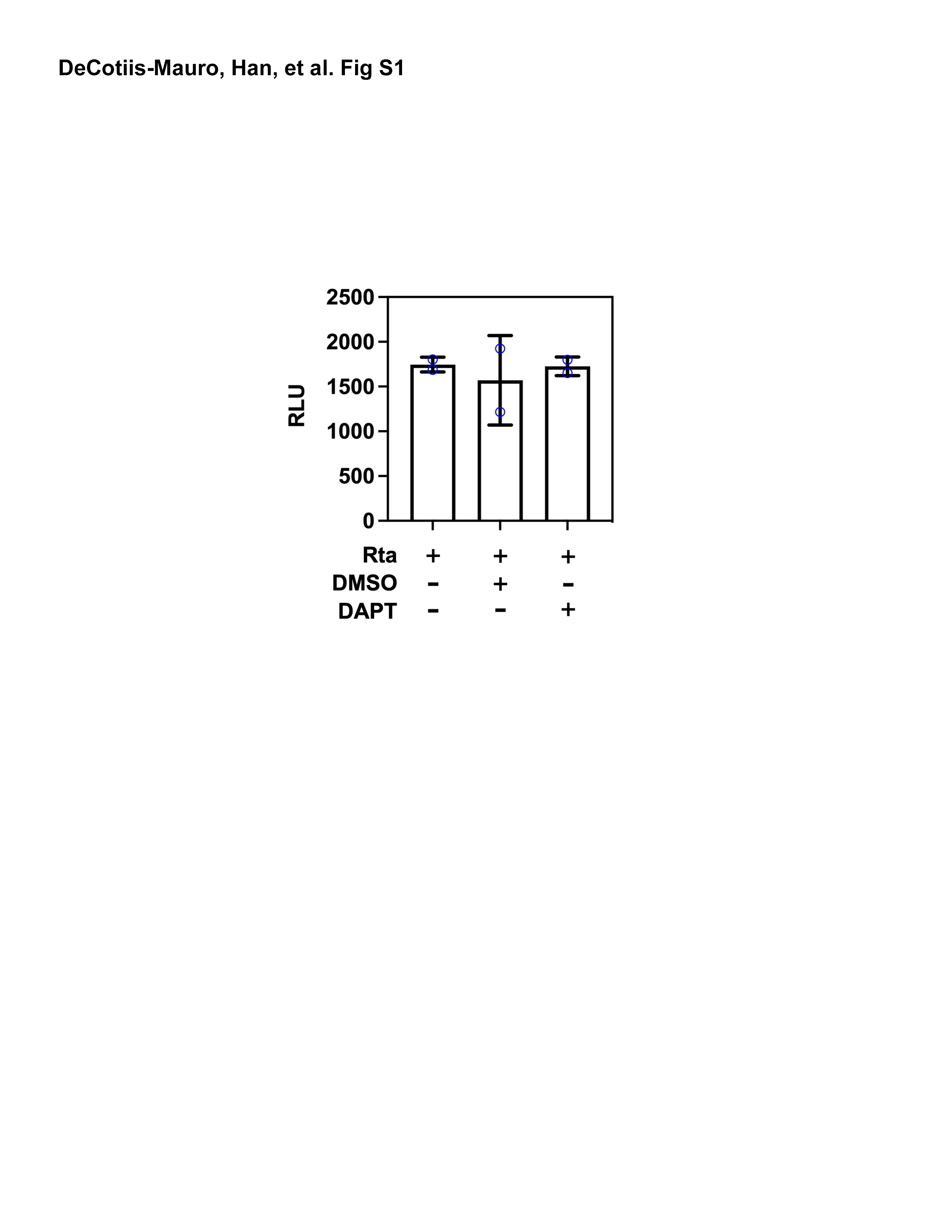

### Supplemental Figure S2

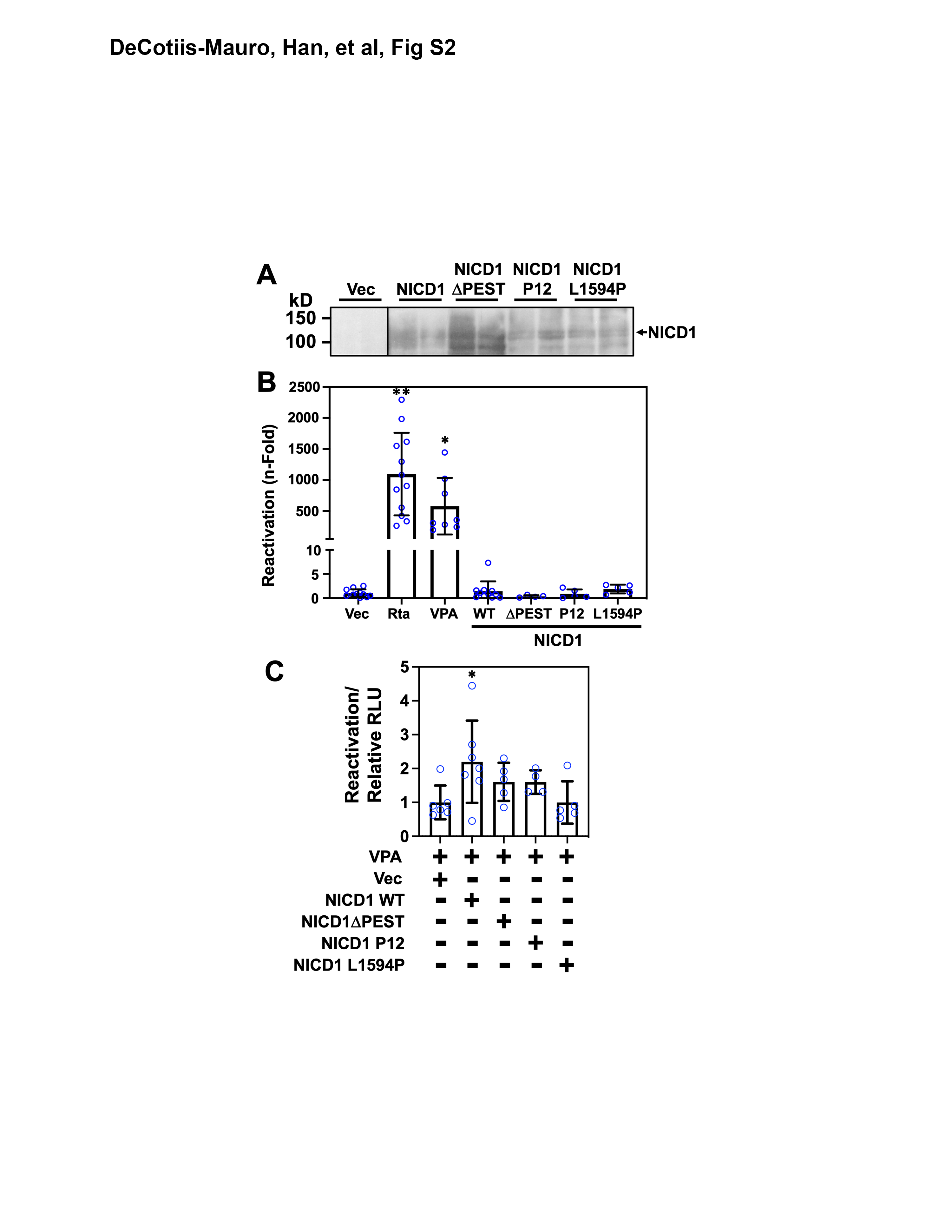
